# Preclinical study of DNA vaccines targeting SARS-CoV-2

**DOI:** 10.1101/2020.10.21.347799

**Authors:** Hiroki Hayashi, Jiao Sun, Yuka Yanagida, Takako Otera, Ritsuko Kubota-Kotetsu, Tatsuo Shioda, Chikako Ono, Yoshiharu Matsuura, Hisashi Arase, Shota Yoshida, Ryo Nakamaru, Ryoko Ide, Akiko Tenma, Sotaro Kawabata, Takako Ehara, Makoto Sakaguchi, Hideki Tomioka, Munehisa Shimamura, Sachiko Okamoto, Yasunori Amaishi, Hideto Chono, Junichi Mineno, Takano Komatsuno, Yoshimi Saito, Hiromi Rakugi, Ryuichi Morishita, Hironori Nakagami

## Abstract

To fight against the worldwide COVID-19 pandemic, the development of an effective and safe vaccine against SARS-CoV-2 is required. As potential pandemic vaccines, DNA/RNA vaccines, viral vector vaccines and protein-based vaccines have been rapidly developed to prevent pandemic spread worldwide. In this study, we designed plasmid DNA vaccine targeting the SARS-CoV-2 Spike glycoprotein (S protein) as pandemic vaccine, and the humoral, cellular, and functional immune responses were characterized to support proceeding to initial human clinical trials. After intramuscular injection of DNA vaccine encoding S protein with alum adjuvant (three times at 2-week intervals), the humoral immunoreaction, as assessed by anti-S protein or anti-receptor-binding domain (RBD) antibody titers, and the cellular immunoreaction, as assessed by antigen-induced IFNγ expression, were up-regulated. In IgG subclass analysis, IgG2b was induced as the main subclass. Based on these analyses, DNA vaccine with alum adjuvant preferentially induced Th1-type T cell polarization. We confirmed the neutralizing action of DNA vaccine-induced antibodies via two different methods, a binding assay of RBD recombinant protein with angiotensin-converting enzyme 2 (ACE2), a receptor of SARS-CoV-2, and pseudovirus assay. Further B cell epitope mapping analysis using a peptide array showed that most vaccine-induced antibodies recognized the S2 and RBD subunits, but not the S1 subunit. In conclusion, DNA vaccine targeting the spike glycoprotein of SARS-CoV-2 might be an effective and safe approach to combat the COVID-19 pandemic.

## Introduction

The pandemic of COVID-19 spread from the reported cluster of pneumonia cases in Wuhan, Hubei Province, in Dec 2019. Patients with COVID-19 present with viral pneumonia caused by severe acute respiratory syndrome-coronavirus 2 (SARS-CoV-2) (World Health Organization (WHO) “Novel Coronavirus-China” WHO, 2020: www.who.int/csr/don/12-january-2020-novel-coronavirus-china/en/). The number of infected people reached 14 million as of July 2020 and is increasing worldwide. The development of an effective and safe vaccine to combat this unprecedented global pandemic is urgently needed. Many pharmaceutical companies and academia are developing vaccines against SARS-CoV-2, including adenovirus-based, DNA or RNA-based, and inactivated vaccines ^1, 2, 3, 4, 5^, mostly targeting the spike glycoprotein of SARS-CoV-2, which is essential for virus entry into cells ^6^.

The advantages of DNA vaccines are that they (1) can be simply and quickly produced by PCR or synthetic methods, (2) can be easily produced at a large scale, (3) are safer than other approaches, such as inactivated virus vaccines, and (4) are more thermostable than other types of vaccines ^7^, according to WHO DNA vaccine guideline. SARS-CoV-2, classified as the species severe acute respiratory syndrome-related coronavirus, is a member of the family of enveloped positive-sense RNA viruses ^8^. The mutation rate of RNA viruses is known to be higher than that of DNA viruses ^9^, suggesting that developed vaccines for SARS-CoV-2 need to be adapted for its mutation. In this study, we developed DNA-based vaccine targeting the SARS-CoV-2 spike glycoprotein. In the guidance by the FDA on vaccines to prevent COVID-19, preclinical studies of a COVID-19 vaccine candidate require the evaluation of humoral, cellular, and functional immune responses to support proceeding to initial human clinical trials. Accordingly, in this study, antibody production measured by an antigen-specific enzyme-linked immunosorbent assay (ELISA) was considered to represent the humoral response, and antigen-dependent T cell activation by an enzyme-linked immunosorbent spot (ELISpot) assay was evaluated for cellular responses. The functional activity of immune responses was evaluated *in vitro* in neutralization assays using pseudovirion virus. The assays used for immunogenicity evaluation should be demonstrated to be suitable for their intended purpose. Furthermore, we conducted B cell epitope analysis for the induced antibodies.

## Results

### DNA vaccine design and in vitro expression

The optimized DNA sequence of the SARS-CoV-2 (isolate Wuhan-Hu-1; MN_908947.3) spike glycoprotein was inserted into the pVAX1 plasmid (Fig. 1a). The spike glycoprotein contains the RBD, heptad repeat 1 (HR1), heptad repeat 2 (HR2), the transmembrane domain, and the cytosolic domain. The expression of the spike glycoprotein was confirmed by western blot. The construct was transfected into HEK293 cells and incubated for 48 hours. The spike glycoprotein was detected in HEK293 cells transfected with the DNA vaccine construct using a specific anti-Spike glycoprotein antibody, and the recombinant protein was almost the same size as the band of recombinant S1+S2 (extracellular domain: ECD, Fig. 1b). To quantify the expression level of the spike glycoprotein in the cells or in the supernatant, a sandwich ELISA was developed by using anti-Spike glycoprotein antibodies. It has been reported that SARS-CoV-2 infects host cells via ACE2 ^6^. We are concerned with the possibility that the spike glycoprotein originating from DNA vaccine could bind to host ACE2 to regulate its function, including cardiopulmonary function ^10, 11^. Thus, we also evaluated the secretion of the spike glycoprotein in the culture medium of HEK293 cells transiently overexpressing the SARS-CoV-2 spike protein. Indeed, the expression of the spike glycoprotein was detected in the lysate of transfected cells, but not in the supernatant of transfected HEK293 cells (Fig. 1c). The localization of expressed spike glycoprotein was also analyzed by immunostaining in HEK293 cells. At 48 hours after transfection, cells were fixed and stained with an anti-spike glycoprotein antibody. The spike protein was localized mainly on the plasma membrane (Fig. 1d and Supplementary Fig. 1). These in vitro studies confirmed the expression of the spike glycoprotein from the DNA vaccine construct.

**Figure 1.**
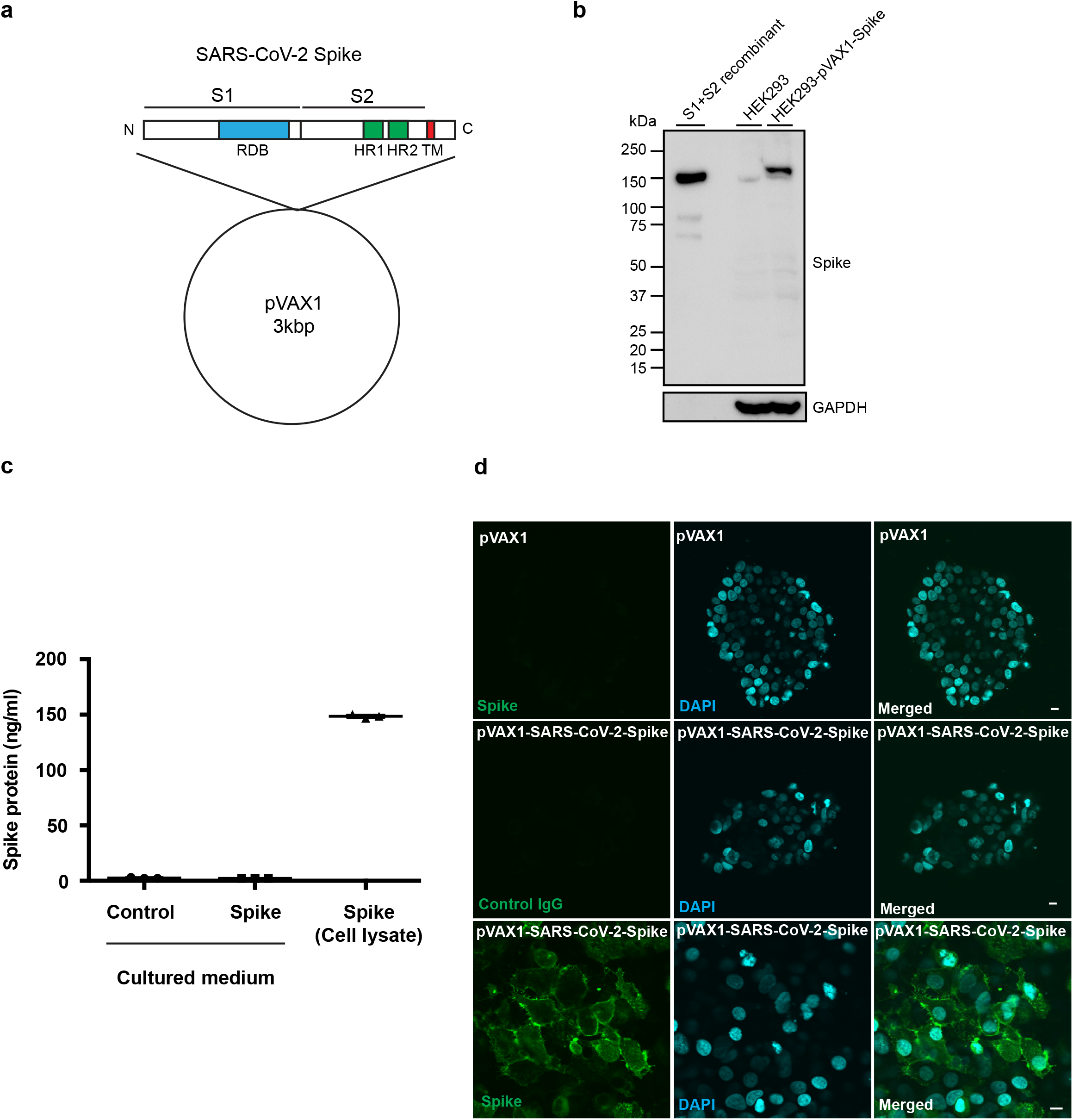
DNA vaccine design and expression of DNA plasmids in cells. (a) Design of DNA vaccine encoding SARS-CoV-2 spike protein based on pVAX1. The Spike glycoprotein is composed of an S1 subunit containing the RBD (receptor-binding domain), an S2 subunit containing HR1 (heptad repeat 1) and HR2 (heptad repeat 2), a transmembrane domain and a cytosolic domain. (b) In vitro expression of pVAX1-SARS-CoV-2 Spike in HEK293 cells as assessed by western blot. Transfected HEK293 cell lysate (HEK293-pVAX-Spike), nontransfected HEK293 cell lysate (HEK293) and recombinant spike glycoprotein (S1+S2 recombinant) were separated by SDS-PAGE and transferred to PVDF membranes. The Spike glycoprotein was detected by the polyclonal anti-spike antibody. GAPDH was used as loading control. (c) Quantification of pVAX1-SARS-CoV-2 Spike protein in HEK293 cells as assessed by sandwich ELISA. The Spike glycoprotein was detected in transfected HEK293 cell lysate (transfected HEK293 lysate), but not in the culture medium of transfected HEK293 cells (sup.). (d) Localization of pVAX1-SARS-CoV-2 Spike glycoprotein in HEK293 cells by immunostaining. The transfected spike protein was stained with the polyclonal spike antibody and the secondary antibody labeled with Alexa Fluor 488 (green). The nucleus was stained with DAPI (blue). Scale bar=10 μm.

### Animal protocol for DNA vaccine administration and evaluating humoral responses in rats

Although plasmid DNA itself induces innate immune responses leading to adjuvant action ^12^, previous reports suggest that the effect of DNA vaccines can be enhanced with alum adjuvant ^13, 14^. To determine whether co-administration of alum adjuvant with DNA vaccine for SARS-CoV-2 enhances antibody production, we compared the DNA vaccine protocol with or without alum adjuvant for 2 weeks. Based on the formulation of several vaccines with alum adjuvants in humans, the dose of alum adjuvant in human clinical trials has been fixed at 0.2 mL (1 mg of aluminum). The compositions of the plasmid DNA and alum adjuvants in this study were calculated based on the clinical protocol (2 mg of DNA plasmid and 1 mg of aluminum). As a result, antibody production was enhanced by alum adjuvant (Fig. 2a), and we selected the co-administration of alum adjuvant for further experiments. The DNA vaccine with alum adjuvant (666.6 μg of plasmid DNA with 66.7 μl of alum adjuvant/rat) was intramuscularly injected into SD rats three times at 2-week intervals (Fig. 2b). DNA vaccine-induced antibody production was followed up to 16 weeks after the 1^st^ vaccination. The antibody titers (half maximum) for the spike glycoprotein and its RBD protein was elevated at 4 through 16 weeks (Fig. 2c, 2d and Supplementary Fig. 2). Moreover, vaccine-induced antibodies also recognized recombinant S1 subunit protein with the D614G mutation of SARS-CoV-2 (Supplementary Fig. 3), which was reported in a European strain, exhibiting strong infectivity compared with that of the Wuhan strain ^15^. At 4 weeks after the 1^st^ vaccination, IgG subclasses (IgG1, IgG2a, IgG2b, and IgG2c) were analyzed by ELISA. Compared with IgG1, IgG2b (and IgG2a) was the main subclass produced by DNA vaccine, suggesting that the humoral immune response shifted toward Th1 response rather than Th2 (Fig. 2e). These data suggest that DNA vaccine effectively activated humoral immune responses.

**Figure 2.**
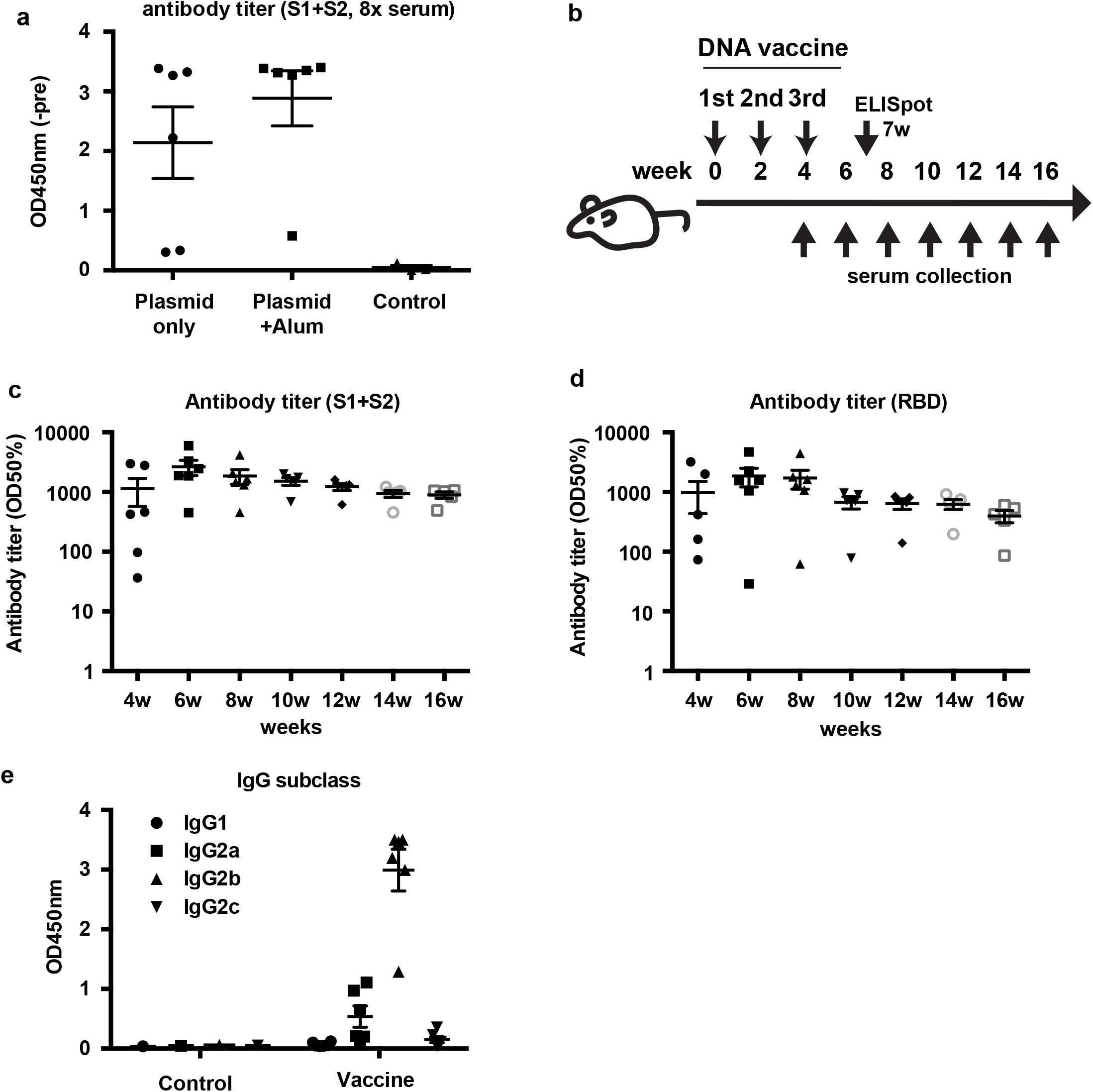
DNA vaccine animal protocol and humoral response induced by the DNA vaccine. (a) Comparison of the effect of DNA vaccine with or without alum adjuvant. Antibody titers were compared at 2 weeks after vaccine administration. (b) Animal protocol for DNA vaccine administration. DNA vaccine was intramuscularly injected with alum adjuvant (666.6 μg of plasmid DNA with 66.7 μl of alum adjuvant/rat) three times at 2-week intervals. Serum samples were collected every two weeks for antibody titer evaluation. (c) Antibody titer for recombinant S1+S2 and (d) antibody titer for recombinant RBD assessed by ELISA from 4 to 16 weeks after the 1^st^ vaccination. Serum dilution from 10x to x31250. The antibody titer is shown as the serum dilution exhibiting half maximum binding at optical density at 450 nm (OD50%). (e) IgG subclasses for recombinant S1+S2 assessed by ELISA at 4 weeks. Serum dilution: 8x dilution.

### The DNA vaccine effectively elicits a cellular immune response in rats

We also examined whether DNA vaccine would induce the cellular immune response by IFNγ and IL4 ELISpot assays at 7 weeks after the 1^st^ vaccination. IFNγ spots were significantly increased by S1+S2 recombinant protein and recombinant RBD protein immunization in rats (Fig. 3a). IL4 spots were slightly increased by S1+S2 recombinant and RBD recombinant protein administration (Fig. 3b). These results and the results of the IgG subclass analysis (Fig. 2e) suggest that DNA vaccine would elicit the cellular immune response toward the Th1 type.

**Figure 3.**
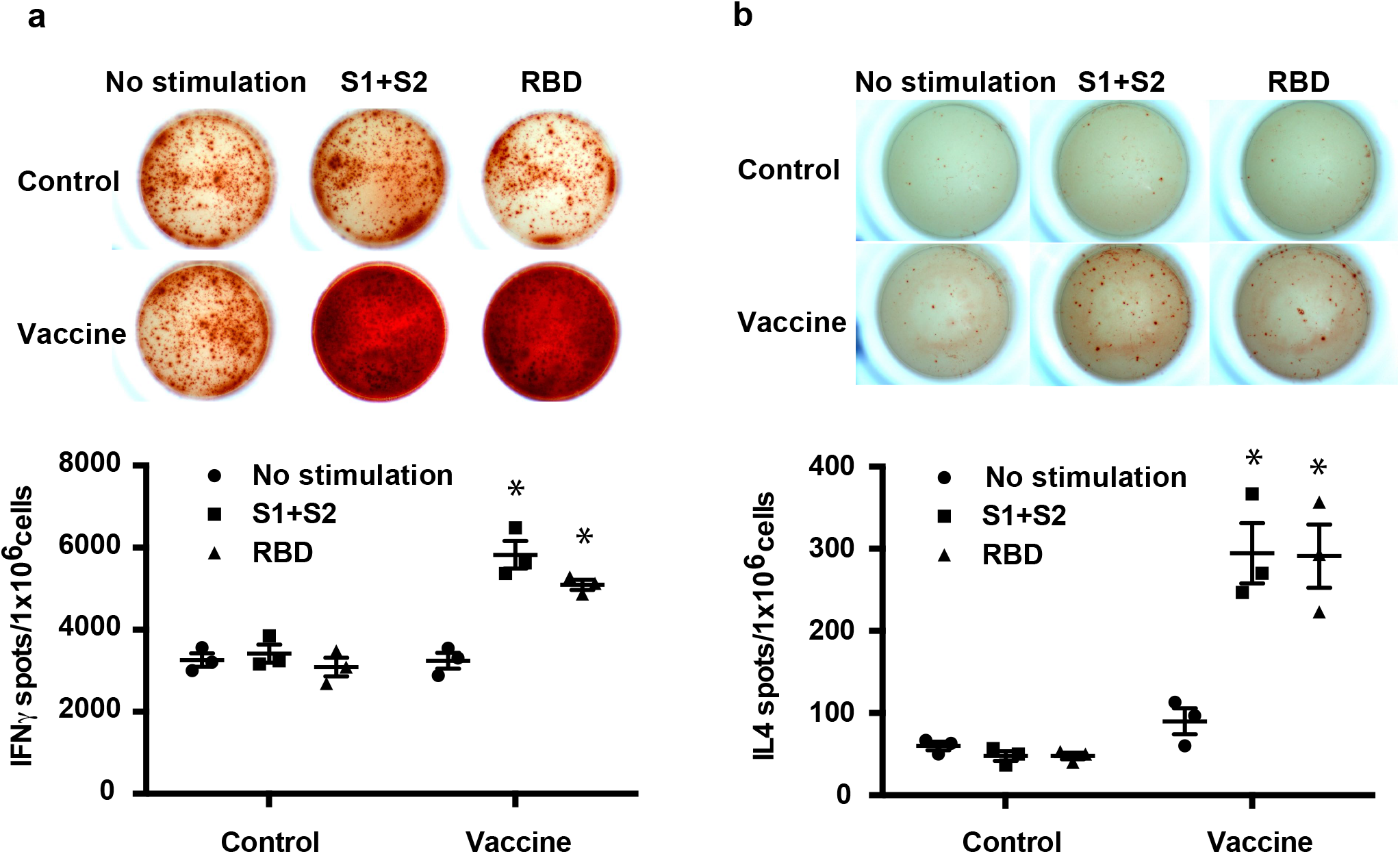
Cellular immune response to the DNA vaccine in rats. (a) IFNγ ELISpot assay and (b) IL4 ELISpot assay in immunized or control rats at 7 weeks after the first vaccination. Splenocytes were stimulated with recombinant S1+S2 or recombinant RBD for 48 hours. *P<0.01 vs. control (splenocytes from untreated rats).

### The DNA vaccine enhanced neutralizing antibodies

We further evaluated the neutralization activity of the vaccine-induced antibodies against SARS-CoV-2 with two different methods. Binding of human ACE2, a receptor of the SARS-CoV-2 spike glycoprotein ^6^, was evaluated by ELISA. The binding of ACE2 with S1+S2 recombinant protein was inhibited by a 5-fold dilution of immunized rat serum, and the binding of ACE2 and RBD protein was also decreased (Fig. 4a). Moreover, neutralizing antibodies were tested by pseudotyped VSV with the luciferase gene and Vero E6 cells stably expressing TMPRSS2 ^6, 16^. A series of dilutions of serum at 8 weeks after the first vaccination exhibited neutralizing activity on pseudovirus infection (Fig. 4b), and neutralizing titers (75% inhibitory dose (ID75) shows the serum dilution that caused a 75 % decrease in RLUs) were 98.4 on average at 8 weeks after the 1^st^ administration of the DNA vaccine (Fig. 4c). We further evaluated the neutralizing activity by utilizing live SARS-CoV-2 in two different method (FRNT and TCID). The neutralizing titers (ID 50: 50% inhibitory dose) in FRNT method were 40.7 on average (Fig. 4d, 4e, and Supplementary Fig. 4), and 10-80 in TCID method (Table 1 and Supplementary Fig. 5) 8 weeks after the 1^st^ administration of the DNA vaccine.

**Figure 4.**
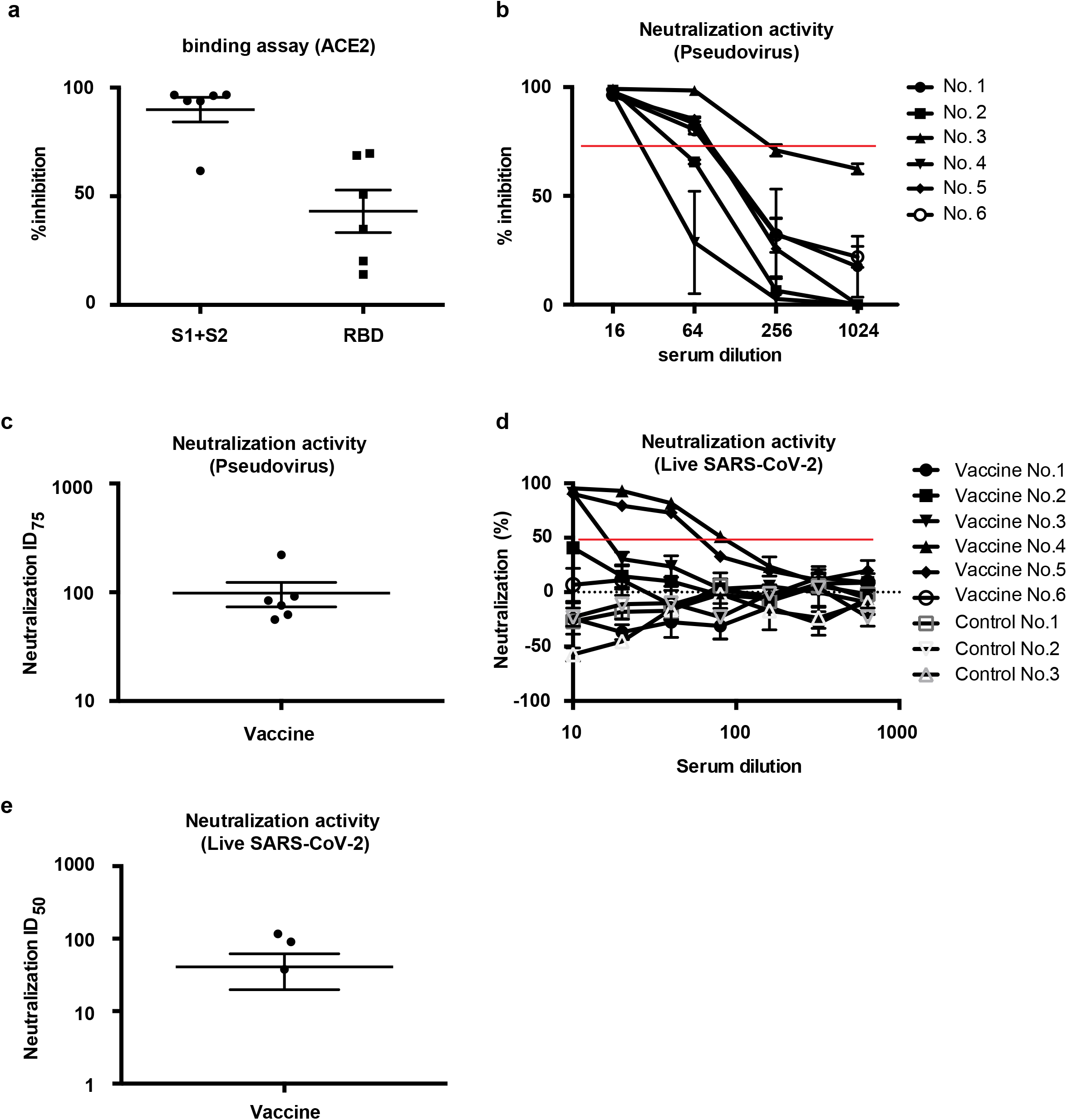
Neutralizing activity of DNA vaccine-induced antibodies. (a) The binding of recombinant ACE2 and recombinant S1+S2 or recombinant RBD assessed by ELISA. The immunized rat serum at 8 weeks was used for ELISA. (b) Neutralization assay using pseudovirus. The immunized rat serum at 8 weeks was used at dilutions of x16, x64, x256, and x1024. (c) The values indicate that the inhibitory dose of serum shows 50 % inhibition. (d) Neutralization assay using live SARS-CoV-2. The immunized rat serum at 8 weeks was used at dilutions of x10, x20, x40, x80, x160, x320, x640. (e) The values indicate that the inhibitory dose of serum shows 50 % inhibition. Vaccine No.3 and Control No.3: 7 weeks sample.

**Table 1.** The result of neutralization activity of vaccinated serum by TCID.

| No. | Neutralization activity |
| --- | --- |
| Vaccine No.1 | <10 |
| Vaccine No.2 | 10 |
| Vaccine No.3 | 80 |
| Vaccine No.4 | 20 |
| Vaccine No.5 | 80 |
| Vaccine No.6 | 10 |
| Control No.1 | <10 |
| Control No.2 | <10 |
| Control No.3 | <10 |

### Evaluation of epitopes in the SARS-CoV-2 spike protein

To identify the epitopes recognized by vaccine-induced antibodies, a peptide array based on the SARS-CoV-2 spike protein was performed with control serum or vaccine serum. The spike peptide-coated membrane treated with immunized rat serum showed many more dots than that treated with control serum (Supplementary Fig. 6). Signals (dots) were ranked according to the strength of the signal for individual samples (Supplementary Table 1), and the top 30 signals were determined from 6 rats (Fig. 5 and Table 2). The top 10 epitopes were localized in the RBD, HR1, HR2, and amino acids approximately 600-700 (near the S1/S2 cleaved site). These data suggest that mainly antibodies recognizing the RBD, HR1, and HR2 were produced by DNA vaccine with the SARS-CoV-2 spike glycoprotein.

**Figure 5.**
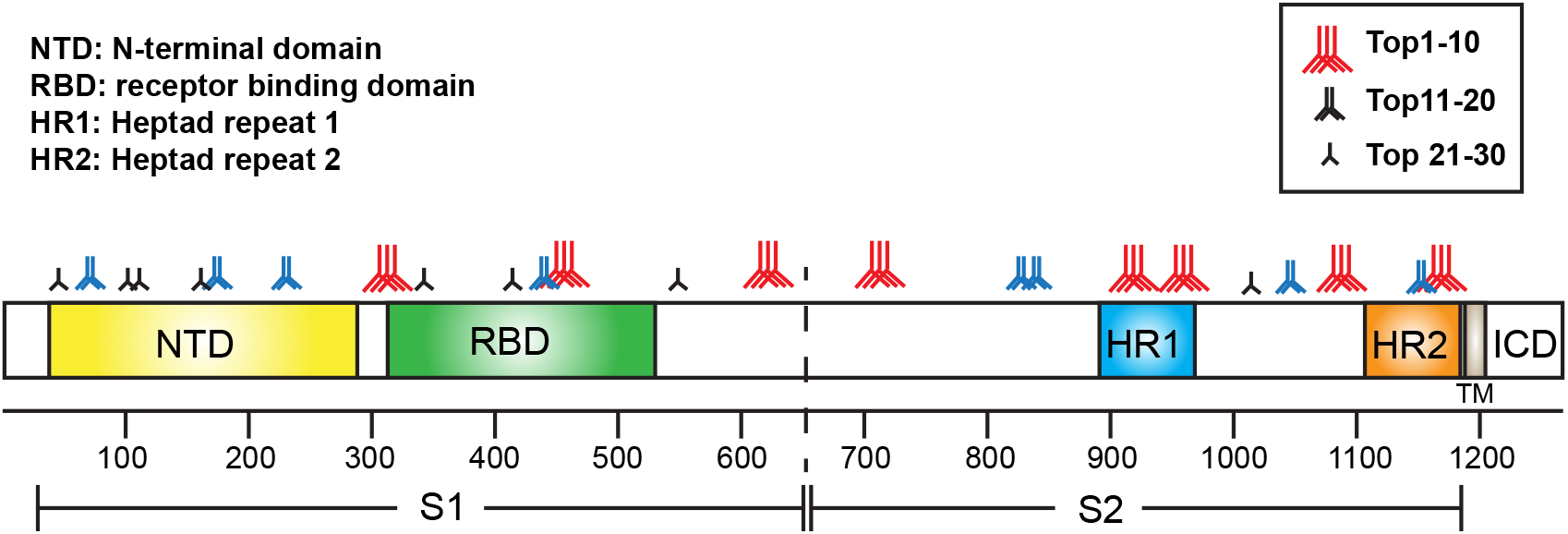
Epitope mapping profiles of DNA vaccine-induced antibodies. The antigenic sites in the spike protein were ranked according to the results of epitope array. “Red three antibodies” indicate the top 10 epitopes. “Blue two antibodies” indicate the top 20 epitopes. “Black antibody” indicates the top 30 epitopes. See the details in Table 2 and Supplementary Table 1.

**Table 2.**
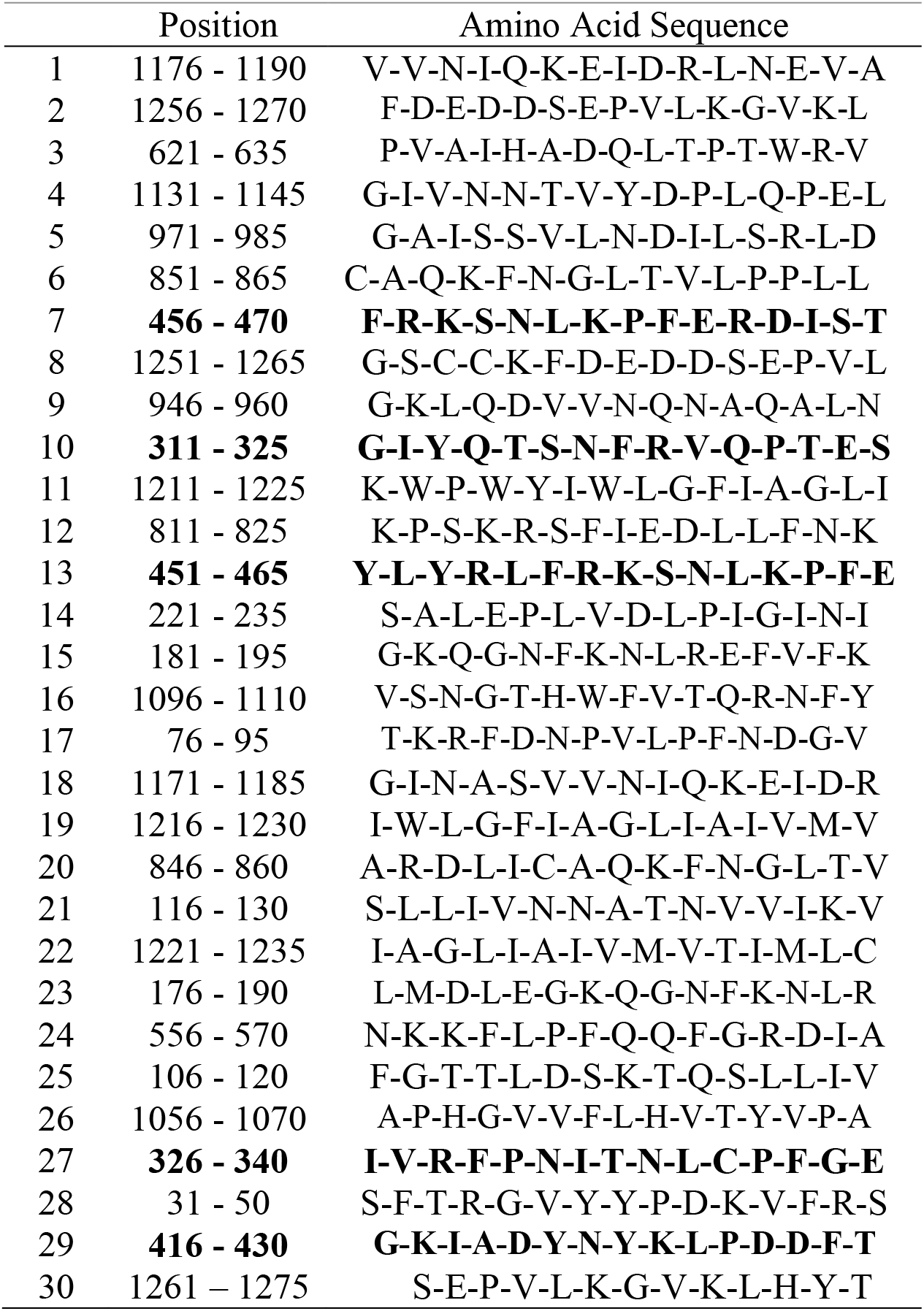
Top 30 strongest epitope recognized by vaccine-induced antibody: bold indicates RBD.

|  | Position | Amino Acid Sequence |
| --- | --- | --- |
| 1 | 1176 - 1190 | V-V-N-I-Q-K-E-I-D-R-L-N-E-V-A |
| 2 | 1256 - 1270 | F-D-E-D-D-S-E-P-V-L-K-G-V-K-L |
| 3 | 621 - 635 | P-V-A-I-H-A-D-Q-L-T-P-T-W-R-V |
| 4 | 1131 - 1145 | G-I-V-N-N-T-V-Y-D-P-L-Q-P-E-L |
| 5 | 971 - 985 | G-A-I-S-S-V-L-N-D-I-L-S-R-L-D |
| 6 | 851 - 865 | C-A-Q-K-F-N-G-L-T-V-L-P-P-L-L |
| 7 | **456 - 470** | **F-R-K-S-N-L-K-P-F-E-R-D-I-S-T** |
| 8 | 1251 - 1265 | G-S-C-C-K-F-D-E-D-D-S-E-P-V-L |
| 9 | 946 - 960 | G-K-L-Q-D-V-V-N-Q-N-A-Q-A-L-N |
| 10 | **311 - 325** | **G-I-Y-Q-T-S-N-F-R-V-Q-P-T-E-S** |
| 11 | 1211 - 1225 | K-W-P-W-Y-I-W-L-G-F-I-A-G-L-I |
| 12 | 811 - 825 | K-P-S-K-R-S-F-I-E-D-L-L-F-N-K |
| 13 | **451 - 465** | **Y-L-Y-R-L-F-R-K-S-N-L-K-P-F-E** |
| 14 | 221 - 235 | S-A-L-E-P-L-V-D-L-P-I-G-I-N-I |
| 15 | 181 - 195 | G-K-Q-G-N-F-K-N-L-R-E-F-V-F-K |
| 16 | 1096 - 1110 | V-S-N-G-T-H-W-F-V-T-Q-R-N-F-Y |
| 17 | 76 - 95 | T-K-R-F-D-N-P-V-L-P-F-N-D-G-V |
| 18 | 1171 - 1185 | G-I-N-A-S-V-V-N-I-Q-K-E-I-D-R |
| 19 | 1216 - 1230 | I-W-L-G-F-I-A-G-L-I-A-I-V-M-V |
| 20 | 846 - 860 | A-R-D-L-I-C-A-Q-K-F-N-G-L-T-V |
| 21 | 116 - 130 | S-L-L-I-V-N-N-A-T-N-V-V-I-K-V |
| 22 | 1221 - 1235 | I-A-G-L-I-A-I-V-M-V-T-I-M-L-C |
| 23 | 176 - 190 | L-M-D-L-E-G-K-Q-G-N-F-K-N-L-R |
| 24 | 556 - 570 | N-K-K-F-L-P-F-Q-Q-F-G-R-D-I-A |
| 25 | 106 - 120 | F-G-T-T-L-D-S-K-T-Q-S-L-L-I-V |
| 26 | 1056 - 1070 | A-P-H-G-V-V-F-L-H-V-T-Y-V-P-A |
| 27 | **326 - 340** | **I-V-R-F-P-N-I-T-N-L-C-P-F-G-E** |
| 28 | 31 - 50 | S-F-T-R-G-V-Y-Y-P-D-K-V-F-R-S |
| 29 | **416 - 430** | **G-K-I-A-D-Y-N-Y-K-L-P-D-D-F-T** |
| 30 | 1261 - 1275 | S-E-P-V-L-K-G-V-K-L-H-Y-T |

### The DNA vaccine has no toxic effect

To evaluate the toxic effect of DNA vaccine on tissues, the lung, liver, kidney, and heart were collected from immunized rats at 7 weeks and analyzed by HE staining. The HE stained sections showed no tissue toxicity (Fig. 6a). Additionally, serum biochemical parameters were evaluated to confirm the tissue toxic effect of the DNA vaccine. Serum biochemical data showed that AMY levels were slightly down-regulated by DNA vaccine compared with those in control serum. However, this variation was within the normal range. Other parameters were not changed (Fig. 6b-o), suggesting that DNA vaccine does not have the toxic effects on tissues.

**Figure 6.**
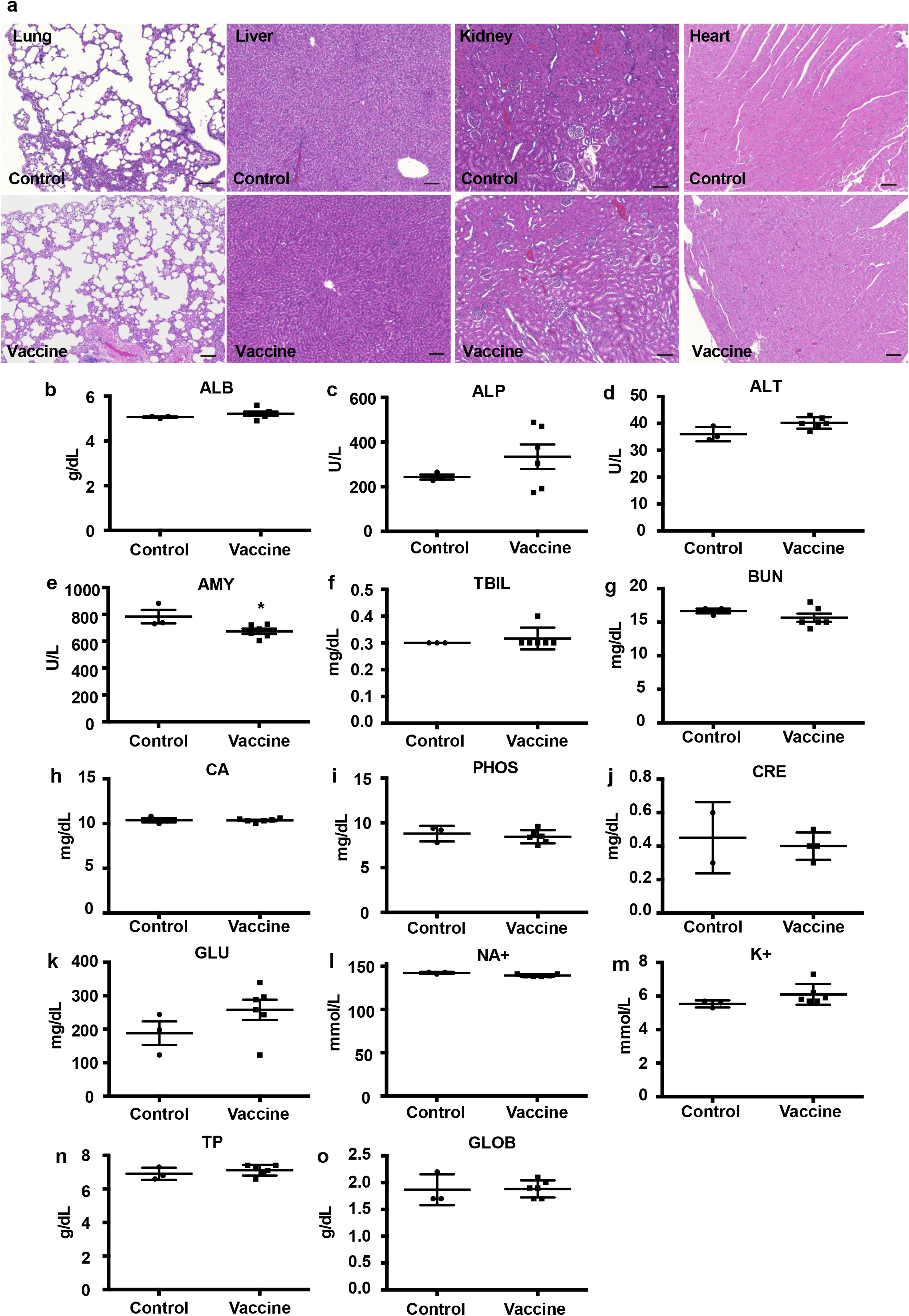
No tissue toxicity with DNA vaccine administration. (a) No tissue toxic effect with DNA vaccine administration assessed by histological analysis (HE staining; lung, liver, kidney, and heart). Scale bar = 100 μm. (b-o) Biochemical analysis of serum at 8 weeks after the 1^st^ vaccination. ALB; albumin, ALP; alkaline phosphatase, ALT; alanine aminotransferase, AMY; amylase, TBIL; total bilirubin, BUN; blood urea nitrogen, CA; calcium, PHOS; phosphate, CRE; creatine, GLU; glucose, NA+; sodium, K+; potassium, TP; total protein, GLOB; globulin. *p<0.05 vs control.

## Discussion

Here, we described the pre-clinical efficacy and safety studies of DNA vaccines for SARS-Cov-2. In this study, we confirmed the expression and immunogenicity of DNA vaccine in *in vitro* and *in vivo* studies, and evaluated the humoral, cellular, and functional immune responses to support proceeding to initial human clinical trials.

DNA vaccine has several potential advantages, including the stimulation of both B and T cell responses, no observed integration of the vector into genomic DNA, rapid construction speed, and good thermostability during storage. DNA vaccine has been applied to many diseases, such as Alzheimer’s disease, allergy, cancer and autoimmune diseases, as well as infections, such as those caused by HIV, hepatitis B, and West Nile virus (WNV) ^17, 18, 19, 20, 21, 22, 23^. For example, DNA vaccine for WNV or Ebola/Marburg viruses efficiently induced immune responses in humans ^24, 25, 26, 27, 28^. In the veterinary field, the protective immune responses have been observed against infectious agents in several target species, including fish, companion animals, and farm animals. DNA vaccines against WNV for use in horses and against infectious hematopoietic necrosis virus (IHNV) for use in salmon were licensed in the USA and in Canada, respectively. DNA vaccine against pancreatic disease was also licensed for use in farmed salmon in several countries ^29^. Toward the clinical application in humans, many approaches have been conducted to enhance the immune response in clinical trials. Although optimization of the plasmid DNA vector (i.e., using strong promoters/enhancers or inserting CpG motifs to enhance adjuvant action) can potentially increase the immunogenicity and strength of gene expression, the plasmid backbone (pVAX) of DNA vaccine was employed in this study, since the pVAX plasmid has already been widely utilized for clinical use with good safety profile, and commercial launched as first gene therapy drug in Japan (Collategene) ^30^. Instead, we optimized the formulation, including co-treatment of polymers, microparticles, utilizing the gene delivery system (intradermal injection, electroporation ^31, 32^), or co-administration of adjuvants ^12, 13, 14^. For the rapid development of DNA vaccine, we selected the co-administration of alum adjuvant that has been clinically used in several vaccines. Although plasmid DNA itself induces innate immune responses leading to adjuvant action ^12^, co-administration of alum adjuvant with DNA vaccine for SARS-CoV-2 enhanced antibody production. The composition of DNA vaccine and alum adjuvants preferentially induced Th1-type T cell polarization based on ELISpot assay and IgG subclass analysis, which might be important to proceed to clinical trials. With model animals administered vaccine constructs against other coronaviruses, evidence of immunopathologic lung reactions characteristic of a Th-2 type hypersensitivity similar to enhanced respiratory disease (ERD) described with the respiratory syncytial virus (RSV) vaccine has been shown ^12, 33, 34, 35, 36, 37^.

*In vivo* animal experiments for SARS-CoV and MERS-CoV vaccines raised serious concerns of the potential risk for SARS-CoV-2 vaccine-associated antibody-dependent enhancement (ADE) ^38, 39^. *In vitro* studies of the effects of antibodies on viral infection have been used extensively to seek the correlates or predictors of ADE. These efforts are complicated by the fact that the antibody mechanisms for protection from viral disease for ADE are similar, and the effect of the administration of passive antibodies has been evaluated for the association with ADE. In small studies, patients infected with SARS-CoV or MERS-CoV received polyclonal antibodies without apparent worsening of their illness, and in the meta-analysis, early treatment with plasma from patients who had recovered from SARS-CoV infection correlated with an improved outcome ^40, 41, 42, 43^. Moreover, a recent meta-analysis found no relationship between the kinetics of antibody responses to SARS-CoV, MERS-CoV or SARS-CoV-2 and clinical outcomes ^44^. The current clinical experience is insufficient to implicate a role for ADE in the severity of COVID-19. To further investigate anti-Spike ADE, B cell epitope analysis was conducted using rat serum including DNA vaccine-induced antibody. Importantly, the antibodies produced by DNA vaccine greatly varied among the rats (Fig. 5). Thus, in the initial phase of clinical trials, B cell epitope analysis might be useful to evaluate the correlates or predictors of ADE with clinical symptoms, which include the magnitude and antigen specificity of antibodies, antibody subclasses and T cell subpopulations. To assess the risk of ADE, there is also value in available animal models for predicting the likelihood of such occurrence in humans. Therefore, post-vaccination animal challenge studies will be required in the near future for vaccine development. Currently, phase 1/2 clinical trial using this DNA vaccine has been tested in Japan. Preliminary data of this clinical trial demonstrated no serious adverse effects (data not shown), similar to the previous data of safety profile using DNA vaccines against various infectious disease.

Overall, these initial results describing the immunogenicity of DNA vaccine targeting S protein for SARS-CoV-2 will provide basal evidence toward to the clinical trials. Development of DNA vaccine against SARS-CoV-2 will lead to provide the novel vaccine for COVID-19 with high safety profile.

## Methods

### Animal protocol

Seven-week-old male Sprague-Dawley (SD) rats were purchased from Clea Japan Inc. (Tokyo, Japan), and housed with free access to food and water in a temperature and light cycle-controlled facility. All experiments were approved by the Ethical Committee for Animal Experiments of the Osaka University Graduate School of Medicine. For immunization, DNA vaccine (666.6 μg with 66.7 μl of alum adjuvant/rat) was intramuscularly injected at 0, 2, and 4 weeks with a needle and syringe. Blood samples were collected every 2 weeks until 16 weeks after the first vaccination. At 7 weeks, the spleen, kidney, lung, and heart were collected for further analysis (spleen for ELISpot assay; kidney, lung, and heart for tissue toxicity).

### DNA vaccine

pVAX1 was obtained from Invitrogen (USA). The virus RNA of SARS-CoV-2 (isolate Wuhan-Hu-1; MN_908947.3) was obtained from the National Institute of Infectious Disease (Tokyo, Japan). A highly optimized DNA sequence encoding the SARS-CoV-2 Spike glycoprotein was created using an in silico gene optimization algorithm to enhance expression and immunogenicity and was inserted into the pVAX1 plasmid. Plasmid DNA was amplified and purified by GIGA prep (Qiagen, USA). Alum phosphate, as an adjuvant, was obtained from InVivogen (USA).

### Cells, DNA transfection, western blotting, and immunostaining

Human embryonic kidney 293 (HEK293) cells were cultured in DMEM containing 10 % FBS plus penicillin/streptomycin and grown at 37 °C in a humidified 5 % CO_2_ incubator.

For western blotting, plasmid DNA (pVAX1-SARS-CoV-2 Spike) was transfected with Lipofectamine 2000 (Invitrogen), according to the manufacturer’s instructions. After 48 hours, the cells were washed and lysed with RIPA buffer (Sigma) containing protease inhibitor cocktail (Roche). After sonication and centrifugation, the supernatant was collected and stored at −80 °C until use. The protein concentration was determined by Bradford assay (Takara), and the cell lysate was mixed with 4x Laemmli loading buffer containing β mercapto-ethanol. The boiled samples were separated on gradient gels (4-20 %, Bio-Rad) and transferred to PVDF membranes. After the membrane was blocked with PBS containing 5 % skim milk for 1 hour at room temperature (RT), the membrane was incubated with an antibody against SARS-CoV-2 Spike (GNX135356, GeneTex, Inc.) at 4 °C overnight. The washed membrane was incubated with a secondary antibody labeled with horseradish peroxidase (HRP) (GE healthcare) for 1 hour at RT. After washing, the membrane was developed with a substrate (Chemi-Lumi One L, Nacalai Tesque). The bands were detected by ChemiDoc™ Touch (Bio-Rad).

For immunostaining, the cells were seeded on glass-bottomed dishes (Matsunami glass) and incubated for 24 hours. At 80 % confluence, plasmid DNA (pVAX1 or pVAX1-SARS-CoV-2 Spike) was transfected with Lipofectamine 2000 (Invitrogen), according to the manufacturer’s instructions. After 48 hours, the cells were fixed with 4 % paraformaldehyde and permealized with 0.2 % Triton X-100. The cells were blocked with 5 % skim milk-PBS for 1 hour at RT and incubated with an antibody against SARS-CoV-2 spike (BLSN-005P, Beta Lifescience) or control IgG (Thermo Fisher) at 4 °C overnight. After washing the cells with PBS, the cells were incubated with a secondary antibody labeled with Alexa Fluor 488 (Molecular Probes) for 1 hour at RT. Nuclei were stained with DAPI. The stained cells were observed by confocal microscopy (FV10i, Olympus).

### Antibody titer determination by ELISA

In this study, we collected serum samples from the tail vein every 2 weeks and evaluated antibody titers by ELISA. Briefly, recombinant 2019-nCoV Spike S1+S2 protein (ECD: Beta Lifescience), recombinant 2019-nCoV Spike protein (RBD; Beta Lifescience), recombinant 2019-nCoV Spike protein (S1; Sino Biological), and recombinant 2019-nCoV Spike (S1-D614G; Sino Biological) (1 μg/ml) were coated on 96-well plates on the first day. On the second day, wells were blocked with blocking buffer (PBS containing 5 % skim milk) for 2 hours at RT. The sera were diluted 10-to 31250-fold in blocking buffer and incubated overnight at 4 °C. The next day, the wells were washed with PBS containing 0.05 % Tween^®^20 (PBS-T) and incubated with HRP-conjugated antibodies (GE healthcare) for 3 hours at RT. After washing with PBS-T, wells were incubated with the peroxidase chromogenic substrate 3,3’-5,5’-tetramethyl benzidine (Sigma-Aldrich) for 30 min at RT, and then the reaction was halted with 0.5 N sulfuric acid. The absorbance of the wells was immediately measured at 450 nm with a microplate reader (Bio-Rad). The value of the half-maximal antibody titer of each sample was calculated from the highest absorbance in the dilution range by using Prism 6 software.

### Sandwich ELISA for detection of secreted spike protein

A diluted capture antibody for the spike protein (GTX632604, GeneTex) was coated on a 96-well plate and incubated at 4 °C overnight. After the plate was blocked with PBS containing 5 % skim milk for 2 hours at RT, diluted recombinant protein (S1+S2, Beta Lifescience) as a standard and samples were loaded and incubated at 4 °C overnight. The washed plate was incubated with a detection antibody (40150-R007, Sino Biological) for 2 hours at RT. After washing with PBS-T, a diluted secondary antibody labeled with HRP (GE healthcare) was added and incubated for 3 hours at RT. After washing with PBS-T, HRP was developed by adding the peroxidase chromogenic substrate 3,3’-5,5’-tetramethyl benzidine (Sigma-Aldrich) for 30 min at RT, and then the reaction was halted with 0.5 N sulfuric acid. The absorbance at 450 nm was measured by a microplate reader (Bio-Rad). The concentration of the spike protein was calculated by a standard curve obtained from recombinant S1+S2 protein.

### ELISpot assay

Cellular immune responses were measured by ELISpot assay, according to the manufacturer’s instructions (UCT Biosciences). Ninety-six-well PVDF membrane-bottomed plates (Merck Millipore) were coated with an anti-rat IFNγ capture antibody or IL4 capture antibody and incubated at 4 °C overnight. After the plates were washed with PBS, the plates were blocked with blocking stock solution (UCT Biosciences) for 2 hours at RT. Splenocytes from immunized or control rats were adjusted to 3×10^5^ well and stimulated with recombinant 2019-nCoV Spike S1+S2 protein (ECD: Beta Lifescience) or recombinant 2019-nCoV-Spike protein (RBD; Beta Lifescience) at 37 °C for 48 hours. The washed plates were incubated with a biotinylated polyclonal antibody specific for rat IFNγ or IL4 for 2 hours at 4 °C. After washing with PBS-T, diluted streptavidin-HRP conjugate was added and incubated for 1 hour at RT. HRP was developed with a substrate solution (AEC coloring system, UCT Biosciences), and then the reaction was stopped by rinsing both sides with demineralized water and air drying at RT in the dark. The colored spots were counted using a dissecting microscope (LMD6500, Leica).

### Epitope array

Epitope mapping of vaccine-induced antibody was performed by using a CelluSpots peptide array (CelluSpots™ COVID19_HullB:98.301, Intavis), according to the manufacturer’s instructions. After blocking the membrane with PBS containing 5 % skim milk for 1 hour at RT, diluted serum samples (1:10) were added and incubated overnight at 4 °C. The next day, the membrane was washed with PBS-T and then incubated with HRP-conjugated anti-Rat IgG (1:1000; GE healthcare) for 1 h at RT. After membrane washing, spots were developed by Chemi-Lumi One L (07889-70, Nacalai Tesque). Signals were detected with a ChemiDoc Touch Imaging System (Bio-Rad) and analyzed with Image Lab software version 6.0.1 (Bio-Rad).

### ACE2 binding assay

For the binding of RBD with ACE2, 96 well plate was coated with RBD recombinant protein (0.5 μg/ml, Fc tag; Sino Biological) at 4 ^o^C overnight. After the plate was blocked with PBS containing 5% skim milk for 2 h at RT, the plate was incubated with rat diluted serum 4 ^o^C overnight. Next day, the plate was washed with PBS containing Tween 20 (0.05%) (PBS-T), and then human ACE2 recombinant protein (0.25 μg/ml, His tag; Sino Biological) was added to the wells and incubated for 2 h at RT. After washing with PBS-T, plate was incubated with anti-his antibody conjugated with HRP (abcam) for 2 h at RT. After washing with PBS-T, wells were incubated with the peroxidase chromogenic substrate 3,3’-5,5’-tetramethyl benzidine (Sigma-Aldrich) for 30 min at room temperature, then reaction was halted with 0.5 N sulfuric acid. Absorbance of wells were immediately measured at 450 nm with a microplate reader (Bio-Rad).

For the binding of S1+S2 with ACE2, the 96 well plate was coated with human ACE2 recombinant protein (1μg/ml, mFc tag; Cell Signaling Technology). The plate was blocked with PBS containing 5% skim milk for 2 h at RT. After blocking, pre-incubated sample of serum with recombinant S1+S2 protein (2 μg/ml, His tag; Beta Lifescience) was added to wells and incubated at 4 ^o^C overnight. After the plate was washed with PBS-T, the wells was incubated with anti-his antibody conjugated with HRP (abcam) for 2 h at RT. After washing with PBS-T, the peroxidase chromogenic substrate 3,3’-5,5’-tetramethyl benzidine (Sigma-Aldrich) was added to wells and incubated for 30 min at RT. After stopped the reaction by adding 0.5 N sulfuric acid. The absorbance at 450 nm was measured by a microplate reader (Bio-rad).

### Pseudovirus neutralization assay for SARS-CoV-2

The neutralizing activity of vaccine-induced antibodies was analyzed with pseudotyped vesicular stomatitis virus (VSVs), as previously described ^45^. Briefly, Vero E6 cells stably expressing TMPRSS2 were seeded on 96-well plates and incubated at 37 °C for 24 hours. Pseudoviruses were incubated with a series of dilutions of inactivated rat serum for 1 hour at 37 °C, and then added to Vero E6 cells. At 24 hours after infection, the cells were lysed with cell culture lysis reagent (Promega), and luciferase activity was measured by a Centro XS^3^ LB 960 (Berthold).

### Live neutralization assay for SARS-CoV-2

The neutralizing activity of vaccine-induced antibodies was analyzed by focus reduction neutralization test (FRNT) and tissue culture infectious dose (TCID) methods. SARS-CoV-2 JPN/TY/WK-521 strain obtained from the National Institute of Infectious Disease (Tokyo, Japan) was used in this study. The FRNT was carried out according to a previously described method with slight modification ^46, 47,^. Briefly, neutralization assay was based on the reduction in focus forming units (FFU) after exposing a given amount of virus to the product to be characterized and comparing with the untreated control. VeroE6/TMPRSS2 cells (4×10^4^ cells/well) were seeded on 96-well plates and incubated at 37° for 24 hours. Serum samples were serially diluted (factor 2 dilutions: 1:10, 1:20, 1:40, 1:80, 1:160, 1:320 and 1:640) and incubated with equal volume of live SARS-CoV-2 (50 FFU/30μl) at 37° for 30 minutes. Aliquots of 30 μl/well of each diluted serum-virus complex were added in VeroE6/TMPRSS2 cells and inoculation for 30 minutes (37◦C; 5% CO_2_). After inoculation, cells were added 50 μl/well of 1% carboxymethyl cellulose and 0.5% BSA supplemented Dulbecco’s Modified Eagle Medium (DMEM) and incubated for 20 hours (37◦C; 5% CO_2_). After incubation, plates were fixed with ethanol and virus infectious cells were stained by peroxidase-anti-peroxidase (PAP) method. As first antibody, 1μg/ml of anti-SARS-CoV/SARS-CoV-2 nucleoprotein monoclonal antibody, HM1054 (EastCoast Bio) was used. The TCID assay was carried out as previously described (Transfusion in press, 2020). Briefly, Serum samples were serially diluted and incubated with equal volume of virus solution (100TCID50/50μl) at 37° for 30 minutes. Aliquots of 50 μl of each diluted serum-virus complex were added in VeroE6/TMPRSS2 cells, seeded in 96-well plates and incubated for 2 days (37◦C; 5% CO_2_). The cells were fixed with 10% formaldehyde (Sigma-Aldrich), and stained with 0.1% methylene blue. Acute and convalescent sera of SARS-CoV-2 infected patients were used as negative and positive controls, respectively. The acute-phase serum showed no neutralization at all while the convalescent serum possessed the significant neutralization activity. The use of patients sera with written informed consent was approved by an Ethics Committee of Research Instituted for Microbial Diseases Osaka University (approval number 31-14-1).

### Histological analysis

For histological analysis, tissues (kidney, liver, lung, and heart) were collected at 7 weeks after the first vaccination and fixed in 10 % neutral buffered formalin. Fixed tissues were embedded in paraffin, cut into 5-μm-thick sections and stained with hematoxylin and eosin (HE staining). Stained tissue sections were observed using a BZ-800Z (Keyence).

### Serum biochemical parameters

Rat serum biochemical parameters were measured by a Vetscan VS2 (Abaxis, Tokyo, Japan) analyzer with a multirotor II VCDP (Abaxis), according to the manufacturer’s instructions. Biochemical parameters, such as albumin (ALB), alkaline phosphatase (ALP), alanine aminotransferase (ALT), amylase (AMY), total bilirubin (TBIL), blood urea nitrogen (BUN), total calcium (CA), phosphorus (PHOS), creatinine (CRE), glucose (GLU), NA^+^, K^+^, total protein (TP), and globulin (GLOB), were analyzed.

### Statistical analysis

All values are presented as the mean±SEM. One-way ANOVA followed by Tukey’s post hoc test was used to assess significant differences in each experiment using Prism 6 software (GraphPad Software). Differences were considered significant when the p value was less than 0.05.

## Acknowledgments

This study was supported by Project Promoting Support for Drug Discovery grants (JP20nk0101602)from the Japan Agency for Medical Research and Development. We thank all members of the Department of Health Development and Medicine, especially, Ms. Satoe Kitabata for secretarial support.

## Author contribution

H.H. and H.N. wrote the manuscript. H.H., T.S., Y.M., H.A., J.M., H.R., R.M. and H.N. designed the study with discussion and revised the manuscript. H.H., J.S., Y.Y., T.O., R.K., C.O., S.Y, R.N., R.I., A.T., S.K., T.E., M.S., H.T., M.S., S.O., Y.A., H.C., T.K. and Y.S. performed the experiments and analyzed the data.

## Competing interests

The Department of Health Development and Medicine is an endowed department supported by Anges, Daicel, and FunPep. The Department of Clinical Gene Therapy is financially supported by Novartis, AnGes, Shionogi, Boeringher, Fancl, Saisei Mirai Clinics, Rohto and Funpep. R.M. is a stockholder of FunPep and Anges. T.O. T.K. and Y.S. are employees of Anges. R.I, A.T, H.K, S.K, E.T, S.M, and H.T are employees of FunPep. R.M, H.T, and A.T. are FunPep stockholders. The funder provided support in the form of salaries for authors but did not have any additional role in the study design, data analysis, decision to publish, or preparation of the manuscript. All other authors declare no competing interests.

## References

1. Corbett KS, et al. SARS-CoV-2 mRNA Vaccine Development Enabled by Prototype Pathogen Preparedness. Preprint at https://www.biorxiv.org/content/10.1101/2020.06.11.145920v1.full.pdf (2020).

2. Gao Q, et al. Development of an inactivated vaccine candidate for SARS-CoV-2. Science 369, 77–81 (2020).

3. Jackson LA, et al. An mRNA Vaccine against SARS-CoV-2 - Preliminary Report. N Engl J Med, (2020).

4. Smith TRF, et al. Immunogenicity of a DNA vaccine candidate for COVID-19. Nat Commun 11, 2601 (2020).

5. Yu J, et al. DNA vaccine protection against SARS-CoV-2 in rhesus macaques. Science, (2020).

6. Hoffmann M, et al. SARS-CoV-2 Cell Entry Depends on ACE2 and TMPRSS2 and Is Blocked by a Clinically Proven Protease Inhibitor. Cell 181, 271–280 e278 (2020).

7. Kutzler MA, Weiner DB. DNA vaccines: ready for prime time? Nat Rev Genet 9, 776–788 (2008).

8. Coronaviridae Study Group of the International Committee on Taxonomy of V. The species Severe acute respiratory syndrome-related coronavirus: classifying 2019-nCoV and naming it SARS-CoV-2. Nat Microbiol 5, 536–544 (2020).

9. Sanjuan R, Nebot MR, Chirico N, Mansky LM, Belshaw R. Viral mutation rates. J Virol 84, 9733–9748 (2010).

10. Crackower MA, et al. Angiotensin-converting enzyme 2 is an essential regulator of heart function. Nature 417, 822–828 (2002).

11. Imai Y, et al. Angiotensin-converting enzyme 2 protects from severe acute lung failure. Nature 436, 112–116 (2005).

12. Ishii KJ, et al. TANK-binding kinase-1 delineates innate and adaptive immune responses to DNA vaccines. Nature 451, 725–729 (2008).

13. Grunwald T, Ulbert S. Improvement of DNA vaccination by adjuvants and sophisticated delivery devices: vaccine-platforms for the battle against infectious diseases. Clin Exp Vaccine Res 4, 1–10 (2015).

14. Wang S, et al. Enhanced type I immune response to a hepatitis B DNA vaccine by formulation with calcium- or aluminum phosphate. Vaccine 18, 1227–1235 (2000).

15. Korber B, et al. Tracking Changes in SARS-CoV-2 Spike: Evidence that D614G Increases Infectivity of the COVID-19 Virus. Cell, (2020).

16. Matsuyama S, et al. Enhanced isolation of SARS-CoV-2 by TMPRSS2-expressing cells. Proc Natl Acad Sci U S A 117, 7001–7003 (2020).

17. Garren H, et al. Phase 2 trial of a DNA vaccine encoding myelin basic protein for multiple sclerosis. Ann Neurol 63, 611–620 (2008).

18. Gottlieb P, Utz PJ, Robinson W, Steinman L. Clinical optimization of antigen specific modulation of type 1 diabetes with the plasmid DNA platform. Clin Immunol 149, 297–306 (2013).

19. Maldonado L, et al. Intramuscular therapeutic vaccination targeting HPV16 induces T cell responses that localize in mucosal lesions. Sci Transl Med 6, 221ra213 (2014).

20. Pierini S, et al. Trial watch: DNA-based vaccines for oncological indications. Oncoimmunology 6, e1398878 (2017).

21. Tebas P, et al. Intradermal SynCon(R) Ebola GP DNA Vaccine Is Temperature Stable and Safely Demonstrates Cellular and Humoral Immunogenicity Advantages in Healthy Volunteers. J Infect Dis 220, 400–410 (2019).

22. Trimble CL, et al. Safety, efficacy, and immunogenicity of VGX-3100, a therapeutic synthetic DNA vaccine targeting human papillomavirus 16 and 18 E6 and E7 proteins for cervical intraepithelial neoplasia 2/3: a randomised, double-blind, placebo-controlled phase 2b trial. Lancet 386, 2078–2088 (2015).

23. Zhu Z, et al. Enhanced Prophylactic and Therapeutic Effects of Polylysine-Modified Ara h 2 DNA Vaccine in a Mouse Model of Peanut Allergy. Int Arch Allergy Immunol 171, 241–250 (2016).

24. Barouch DH, et al. Plasmid chemokines and colony-stimulating factors enhance the immunogenicity of DNA priming-viral vector boosting human immunodeficiency virus type 1 vaccines. J Virol 77, 8729–8735 (2003).

25. Ledgerwood JE, et al. A West Nile virus DNA vaccine utilizing a modified promoter induces neutralizing antibody in younger and older healthy adults in a phase I clinical trial. J Infect Dis 203, 1396–1404 (2011).

26. Martin JE, et al. A West Nile virus DNA vaccine induces neutralizing antibody in healthy adults during a phase 1 clinical trial. J Infect Dis 196, 1732–1740 (2007).

27. Martin JE, et al. A DNA vaccine for Ebola virus is safe and immunogenic in a phase I clinical trial. Clin Vaccine Immunol 13, 1267–1277 (2006).

28. Sarwar UN, et al. Safety and immunogenicity of DNA vaccines encoding Ebolavirus and Marburgvirus wild-type glycoproteins in a phase I clinical trial. J Infect Dis 211, 549–557 (2015).

29. Stenler S, Blomberg P, Smith CI. Safety and efficacy of DNA vaccines: plasmids vs. minicircles. Hum Vaccin Immunother 10, 1306–1308 (2014).

30. Suda H, Murakami A, Kaga T, Tomioka H, Morishita R. Beperminogene perplasmid for the treatment of critical limb ischemia. Expert Rev Cardiovasc Ther 12, 1145–1156 (2014).

31. Jiang J, et al. Integration of needle-free jet injection with advanced electroporation delivery enhances the magnitude, kinetics, and persistence of engineered DNA vaccine induced immune responses. Vaccine 37, 3832–3839 (2019).

32. Schommer NN, et al. Active Immunoprophylaxis and Vaccine Augmentations Mediated by a Novel Plasmid DNA Formulation. Hum Gene Ther 30, 523–533 (2019).

33. Williams JA. Improving DNA vaccine performance through vector design. Curr Gene Ther 14, 170–189 (2014).

34. Li L, Petrovsky N. Molecular mechanisms for enhanced DNA vaccine immunogenicity. Expert Rev Vaccines 15, 313–329 (2016).

35. Marc MA, Dominguez-Alvarez E, Gamazo C. Nucleic acid vaccination strategies against infectious diseases. Expert Opin Drug Deliv 12, 1851–1865 (2015).

36. Wahren B, et al, DNA vaccine: Recent developments and the future. Vaccines. 2,, 785–796 (2014).

37. Tregoning JS, Kinnear E. Using Plasmids as DNA Vaccines for Infectious Diseases. Microbiol Spectr 2, (2014).

38. Iwasaki A, Yang Y. The potential danger of suboptimal antibody responses in COVID-19. Nat Rev Immunol 20, 339–341 (2020).

39. Wang SF, et al. Antibody-dependent SARS coronavirus infection is mediated by antibodies against spike proteins. Biochem Biophys Res Commun 451, 208–214 (2014).

40. Cheng Y, et al. Use of convalescent plasma therapy in SARS patients in Hong Kong. Eur J Clin Microbiol Infect Dis 24, 44–46 (2005).

41. Duan K, et al. Effectiveness of convalescent plasma therapy in severe COVID-19 patients. Proc Natl Acad Sci U S A 117, 9490–9496 (2020).

42. Mair-Jenkins J, et al. The effectiveness of convalescent plasma and hyperimmune immunoglobulin for the treatment of severe acute respiratory infections of viral etiology: a systematic review and exploratory meta-analysis. J Infect Dis 211, 80–90 (2015).

43. Yeh KM, et al. Experience of using convalescent plasma for severe acute respiratory syndrome among healthcare workers in a Taiwan hospital. J Antimicrob Chemother 56, 919–922 (2005).

44. Huang AT, et al, A systematic review of antibody mediated immunity to coronaviruses: antibody kinetics, correlates of protection, and association of antibody responses with severity of disease. Preprint at https://www.medrxiv.org/content/10.1101/2020.04.14.20065771v1.full.pdf (2020).

45. Yoshida S, et al. SARS-CoV-2-induced humoral immunity through B cell epitope analysis and neutralizing activity in COVID-19 infected individuals in Japan. Preprint at https://www.biorxiv.org/content/10.1101/2020.07.22.212761v1.full.pdf (2020).

46. Okuno Y, Tanaka K, Baba K, Maeda A, Kunita N, Ueda S. Rapid focus reduction neutralization test of influenza A and B viruses in microtiter system. J Clin Microbiol 28, 1308–1313 (1990).

47. Ohshima N, Kubota-Koketsu R, Iba Y, Okuno Y, Kurosawa Y. Two types of antibodies are induced by vaccination with A/California/2009 pdm virus: binding near the sialic acid-binding pocket and neutralizing both H1N1 and H5N1 viruses. PLoS One 9, e87305 (2014).

