## Supplement for "Preclinical study of DNA vaccines targeting SARS-CoV-2"

**Supplementrary Figure 1**

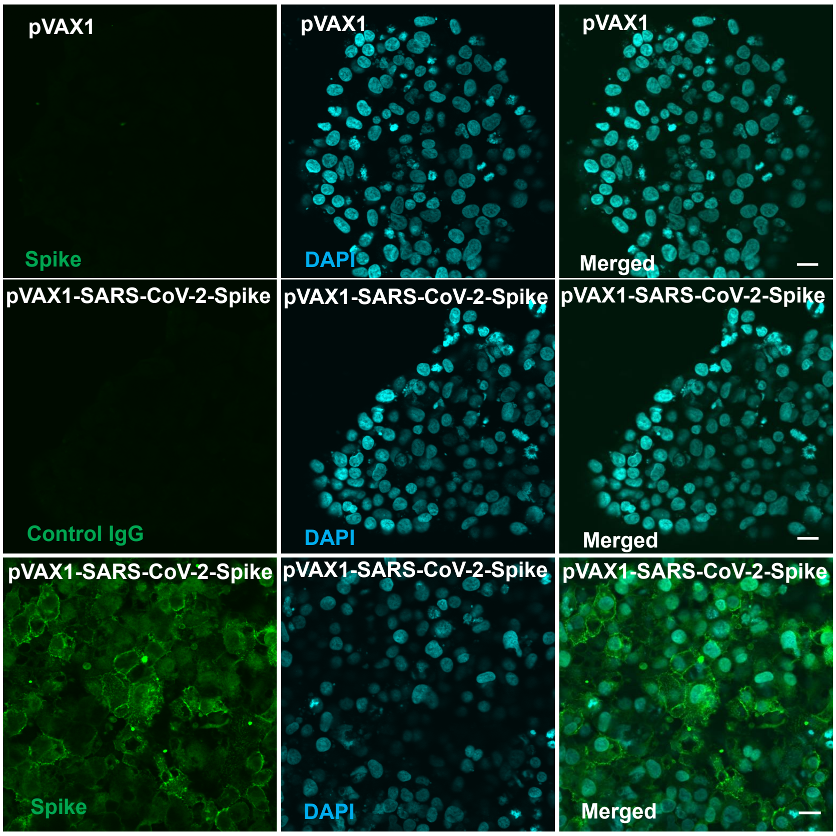


**Supplementary Figure 1. The membrane localization of the spike glycoprotein.** The localization of pVAX1-SARS-CoV-2 Spike in HEK293 cells assessed by immunostaining. The membrane was permeabilized by Triton X-100. The transfected spike protein was stained with a polyclonal spike antibody and a secondary antibody labeled with Alexa Fluor 488 (green). The nucleus was stained with DAPI (blue). Scale bar=20 μm.

**Supplementrary Figure 2**

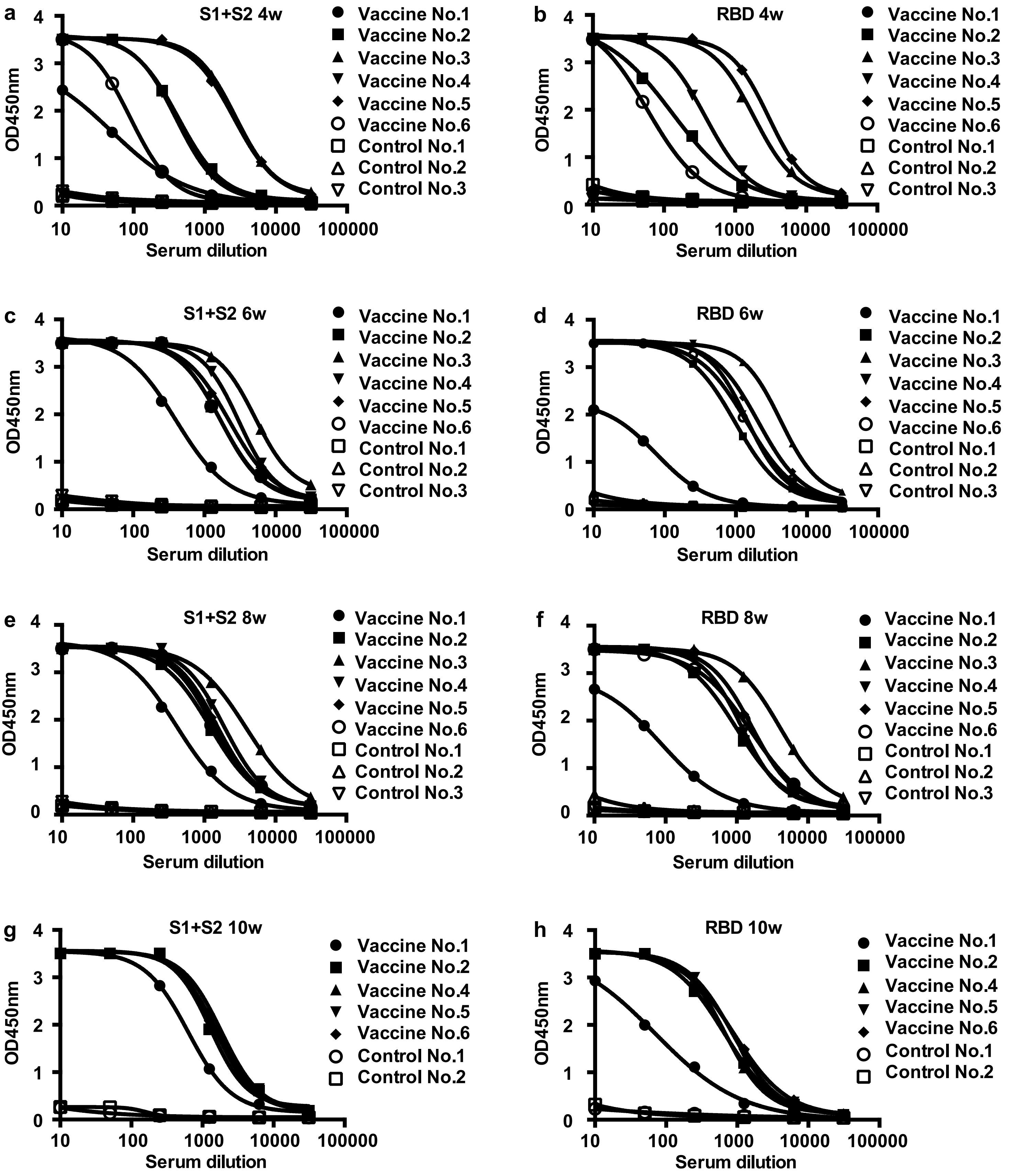

**Supplementrary Figure 2. Continued**

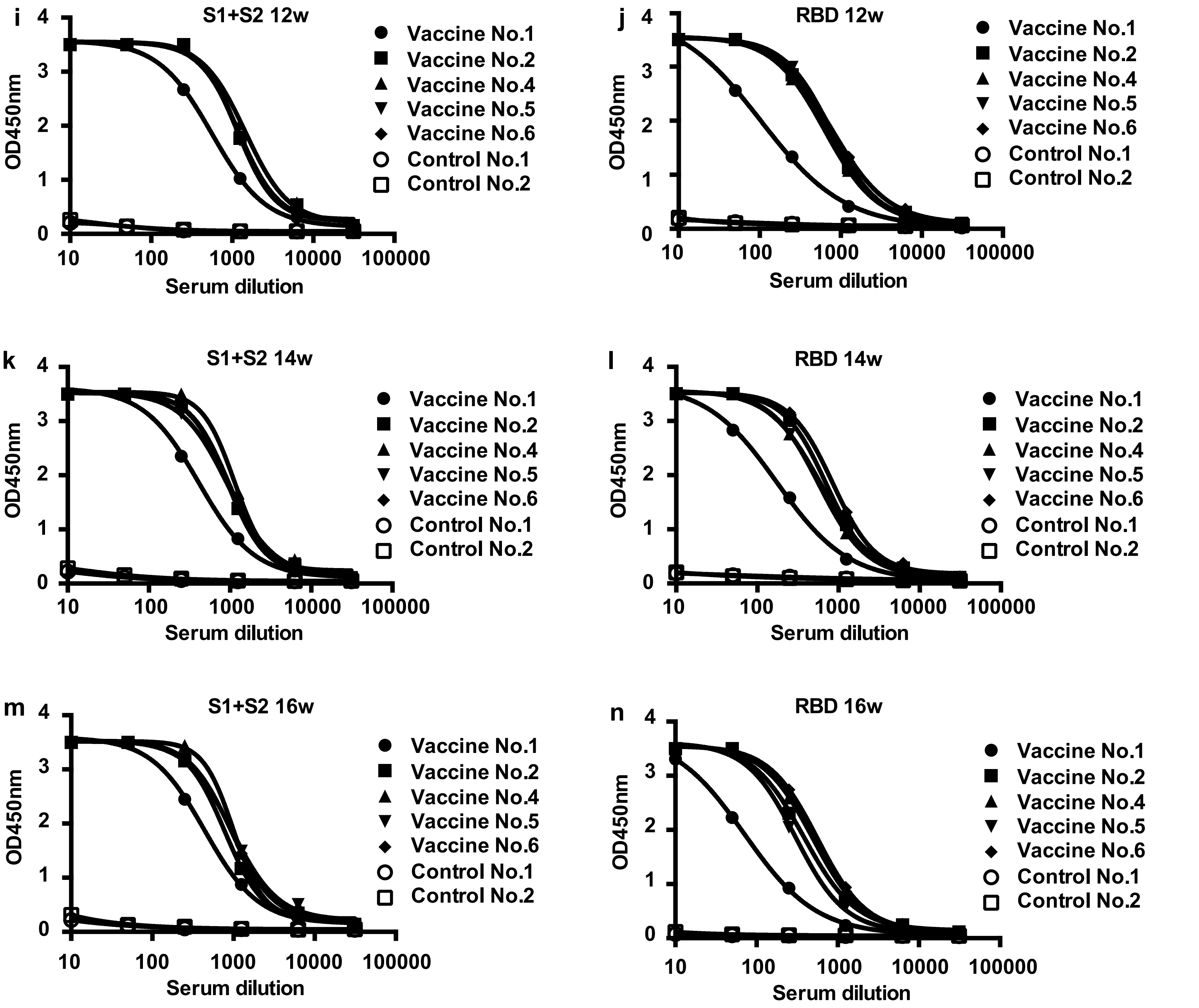


**Supplementary Figure 2. Spike glycoprotein-specific antibody titer from 4 weeks through 16 weeks.** Antibody titers for recombinant S1+S2 and RBD protein measured by ELISA. Serum dilution from 10x to x31250. (**a**) Four weeks: S1+S2. (**b**) Four weeks: RBD. (**c**) Six weeks: S1+S2. (**d**) Six weeks: RBD. (**e**) Eight weeks: S1+S2. Vaccine No.3 and Control No.3: 7 weeks sample. (**f**) Eight weeks: RBD. Vaccine No.3 and Control No.3: 7 weeks sample. (**g**) Ten weeks: S1+S2. (**h**) Ten weeks: RBD. (**i**) Twelve weeks: S1+S2. (**j**) Twelve weeks: RBD. (**k**) Fourteen weeks : S1+S2. (**l**) Fourteen weeks: RBD. (**m**) sixteen weeks : S1+S2. (**n**) sixteen weeks: RBD.

**Supplementary Figure 3**

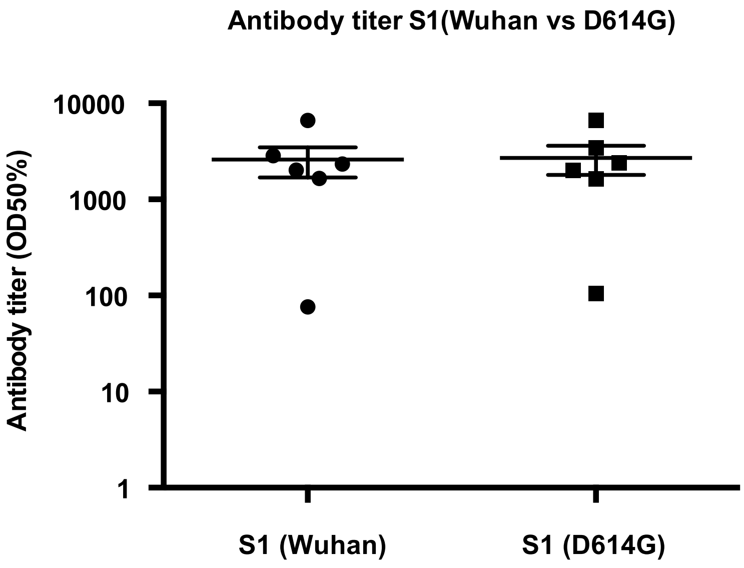


**Supplementary Figure 3. Antibody titer for the S1 subunit with the D614G mutant.** Antibody titer for recombinant S1 subunit from the Wuhan strain or with the D614G mutation assessed by ELISA at 8 weeks after the 1st vaccination. Serum dilution from 10x to x31250. The antibody titer is shown as the serum dilution exhibiting half maximum binding at optical density at 450 nm (OD50%). Vaccine No.3 and Control No.3: 7 weeks sample.

**Supplementary Figure 4**

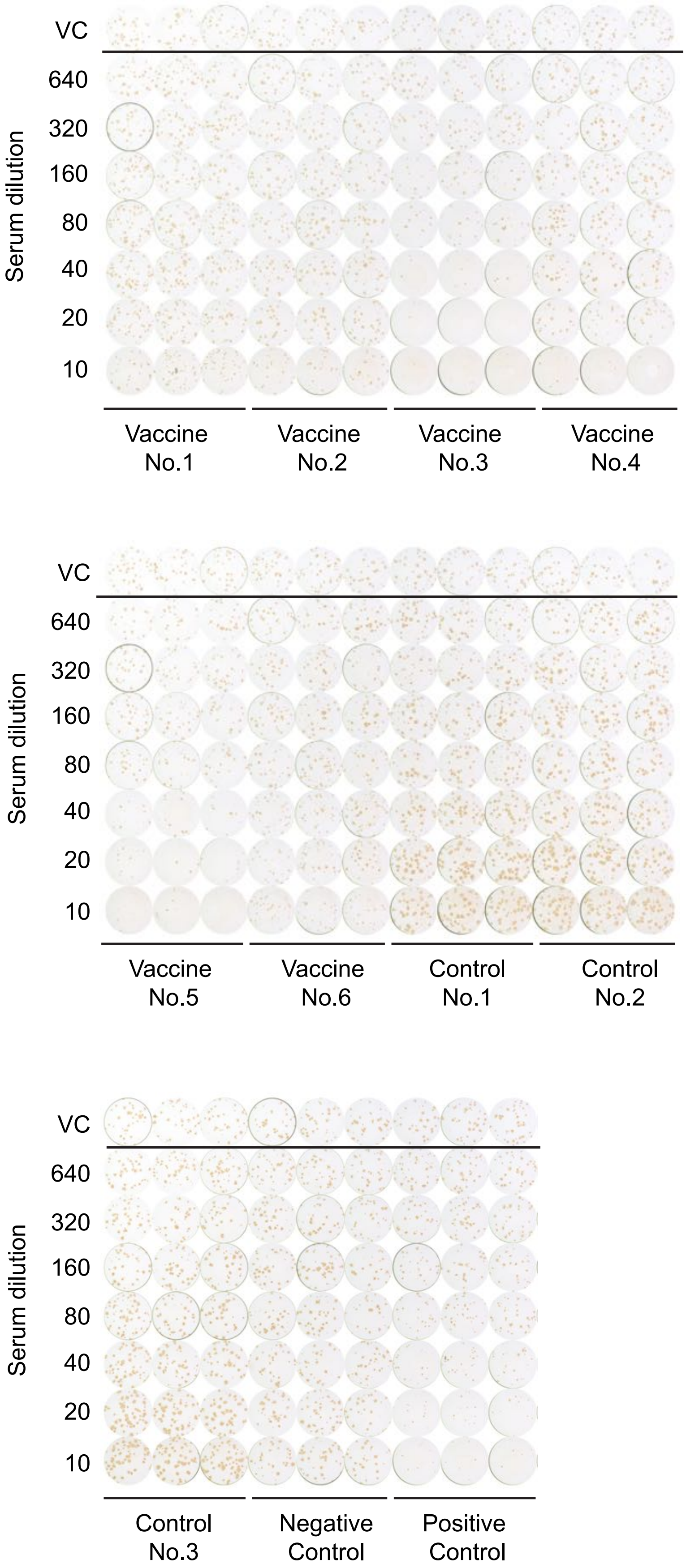


**Supplementary Figure 4. Neutralization activity of vaccinated serum against live SARS-CoV-2 by FRNT.** Neutralization activity of serial dilution of vaccinated or control serum at 8 w was performed with live SARS-CoV-2 infection. VC: viral control. Acute-phase sera of SARS-CoV-2 infected patients was used as negative control, convalescent-phase sera serum of SARS-CoV-2 infected patients was used as positive control.Vaccine No.3 and Control No.3: 7 weeks sample.

**Supplementary Figure 5**

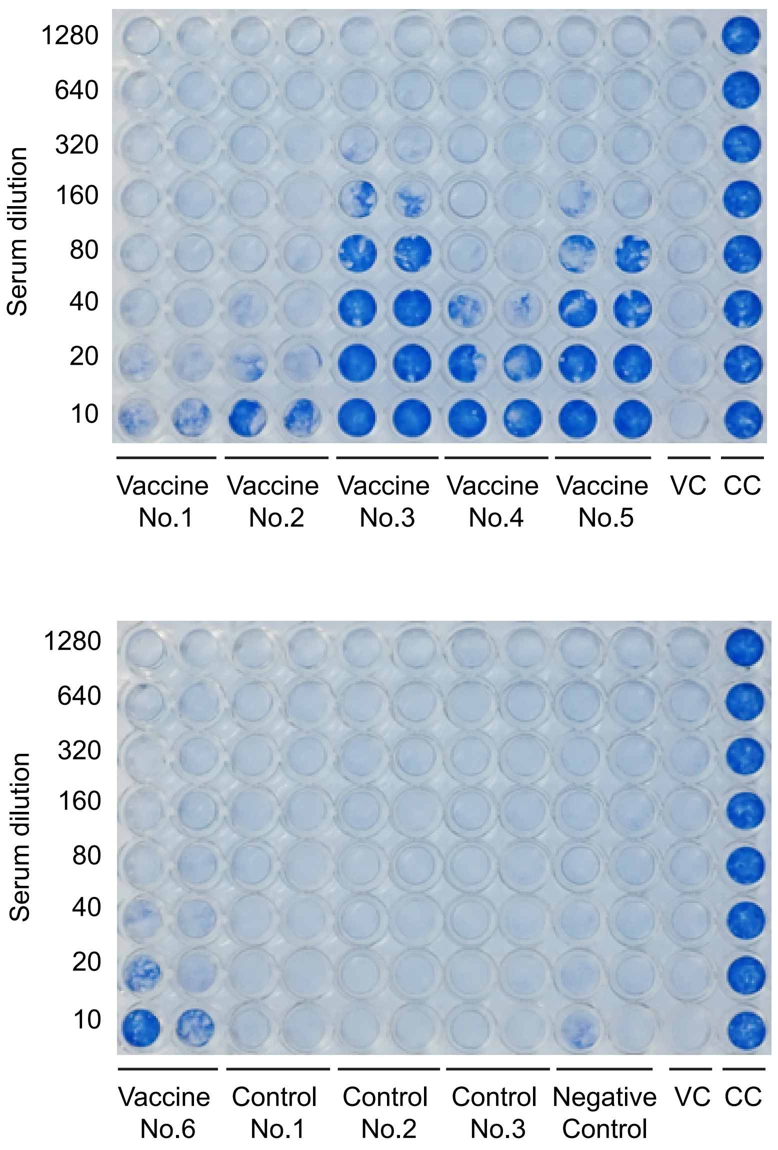


**Supplementary Figure 5. Neutralization activity of vaccinated serum against live SARS-CoV-2 by TCID.** Neutralization activity of serial dilution of vaccinated or control serum at 8w was performed with live SARS-CoV-2 infection. VC: viral control. CC: cell control. Acute-phase sera of SARS-CoV-2 infected patients was used as negative control. Vaccine No.3 and Control No.3: 7 weeks sample.

**Supplementary Figure 6**

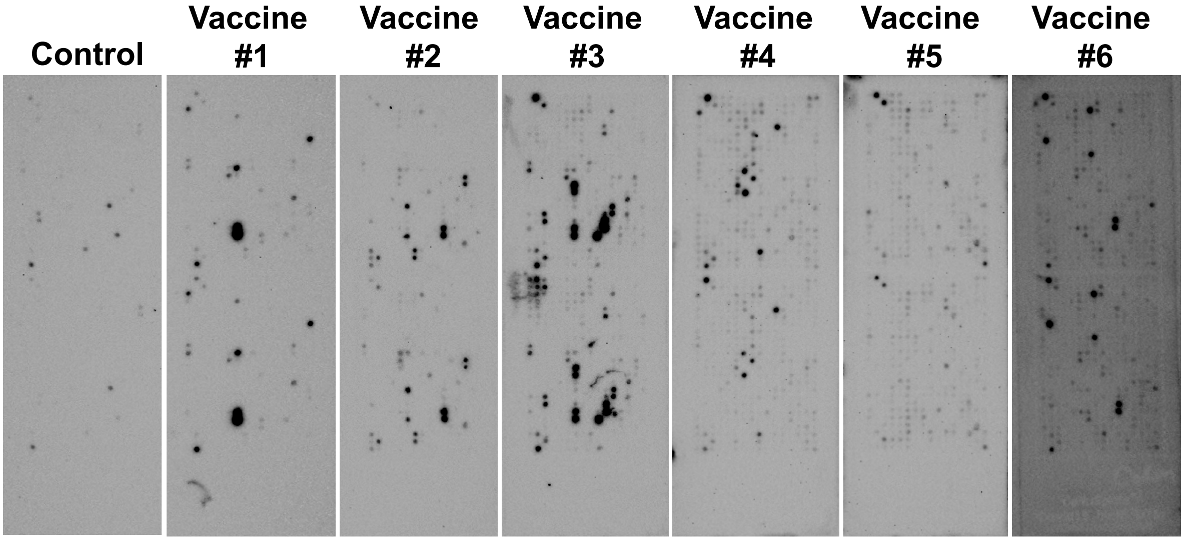


**Supplementary Figure 6. Epitope profiles of vaccine-induced antibodies (peptide array coated with the SARS-CoV-2 Spike glycoprotein).** Spike peptide-coated membranes treated with immunized or nonimmunized rat serum at 6 weeks were developed by chemiluminescence. Images were acquired by a Bio-Rad instrument.

**Supplementary Table 1. Top 30 strongest epitope recognized by vaccine-induced antibody from each animal (No.1-No.6) : bold indicates RBD.**

| No.1 | Position | Amino Acid Sequence |
|---|---|---|
| 1 | 1176 - 1190 | V-V-N-I-Q-K-E-I-D-R-L-N-E-V-A |
| 2 | 1171 - 1185 | G-I-N-A-S-V-V-N-I-Q-K-E-I-D-R |
| 3 | 1131 - 1145 | G-I-V-N-N-T-V-Y-D-P-L-Q-P-E-L |
| 4 | 31 - 50 | S-F-T-R-G-V-Y-Y-P-D-K-V-F-R-S |
| 5 | 1056 - 1070 | A-P-H-G-V-V-F-L-H-V-T-Y-V-P-A |
| 6 | 1166 - 1180 | T-Y-V-P-A-Q-E-K-N-F-T-T-A-P-A |
| 7 | **311 - 325** | **G-I-Y-Q-T-S-N-F-R-V-Q-P-T-E-S** |
| 8 | 1256 - 1270 | F-D-E-D-D-S-E-P-V-L-K-G-V-K-L |
| 9 | 1181 - 1195 | K-E-I-D-R-L-N-E-V-A-K-N-L-N-E |
| 10 | 1051 - 1065 | S-F-P-Q-S-A-P-H-G-V-V-F-L-H-V |
| 11 | 1096 - 1110 | V-S-N-G-T-H-W-F-V-T-Q-R-N-F-Y |
| 12 | 816 - 830 | S-F-I-E-D-L-L-F-N-K-V-T-L-A-D |
| 13 | 286 - 300 | T-D-A-V-D-C-A-L-D-P-L-S-E-T-K |
| 14 | 1136 - 1150 | Q-S-K-R-V-D-F-C-G-K-G-Y-H-L-M |
| 15 | 1251 - 1265 | G-S-C-C-K-F-D-E-D-D-S-E-P-V-L |
| 16 | **456 - 470** | **F-R-K-S-N-L-K-P-F-E-R-D-I-S-T** |
| 17 | 821 - 835 | L-L-F-N-K-V-T-L-A-D-A-G-F-I-K |
| 18 | 81 - 95 | N-P-V-L-P-F-N-D-G-V-Y-F-A-S-T |
| 19 | 1066 - 1080 | T-Y-V-P-A-Q-E-K-N-F-T-T-A-P-A |
| 20 | 36 - 55 | V-Y-Y-P-D-K-V-F-R-S-S-V-L-H-S |
| 21 | 151 - 165 | S-W-M-E-S-E-F-R-V-Y-S-S-A-N-N |
| 22 | 1061 - 1075 | V-F-L-H-V-T-Y-V-P-A-Q-E-K-N-F |
| 23 | 786 - 800 | K-Q-I-Y-K-T-P-P-I-K-D-F-G-G-F |
| 24 | 106 - 120 | F-G-T-T-L-D-S-K-T-Q-S-L-L-I-V |
| 25 | 891 - 905 | G-A-A-L-Q-I-P-F-A-M-Q-M-A-Y-R |
| 26 | 26 - 40 | P-A-Y-T-N-S-F-T-R-G-V-Y-Y-P-D |
| 27 | 291 - 305 | C-A-L-D-P-L-S-E-T-K-C-T-L-K-S |
| 28 | 76 - 95 | T-K-R-F-D-N-P-V-L-P-F-N-D-G-V |
| 29 | **331 - 345** | **N-I-T-N-L-C-P-F-G-E-V-F-N-A-T** |
| 30 | 1126 - 1040 | A-T-K-M-S-E-C-V-L-G-Q-S-K-R-V |

| No.2 | Position | Amino Acid Sequence |
|---|---|---|
| 1 | 576 - 590 | V-R-D-P-Q-T-L-E-I-L-D-I-T-P-C |
| 2 | 571 - 585 | D-T-T-D-A-V-R-D-P-Q-T-L-E-I-L |
| 3 | 176 - 190 | L-M-D-L-E-G-K-Q-G-N-F-K-N-L-R |
| 4 | 1156 - 1170 | F-K-N-H-T-S-P-D-V-D-L-G-D-I-S |
| 5 | 181 - 195 | G-K-Q-G-N-F-K-N-L-R-E-F-V-F-K |
| 6 | 1176 - 1190 | V-V-N-I-Q-K-E-I-D-R-L-N-E-V-A |
| 7 | 1066 - 1080 | T-Y-V-P-A-Q-E-K-N-F-T-T-A-P-A |
| 8 | 221 - 235 | S-A-L-E-P-L-V-D-L-P-I-G-I-N-I |
| 9 | 1071 - 1085 | Q-E-K-N-F-T-T-A-P-A-I-C-H-D-G |
| 10 | 1251 - 1265 | G-S-C-C-K-F-D-E-D-D-S-E-P-V-L |
| 11 | 1256 - 1270 | F-D-E-D-D-S-E-P-V-L-K-G-V-K-L |
| 12 | 566 - 580 | N-K-K-F-L-P-F-Q-Q-F-G-R-D-I-A |
| 13 | 851 - 865 | C-A-Q-K-F-N-G-L-T-V-L-P-P-L-L |
| 14 | **456 - 470** | **F-R-K-S-N-L-K-P-F-E-R-D-I-S-T** |
| 15 | 786 - 800 | K-Q-I-Y-K-T-P-P-I-K-D-F-G-G-F |
| 16 | 1131 - 1145 | G-I-V-N-N-T-V-Y-D-P-L-Q-P-E-L |
| 17 | **311 - 325** | **G-I-Y-Q-T-S-N-F-R-V-Q-P-T-E-S** |
| 18 | **451 - 465** | **Y-L-Y-R-L-F-R-K-S-N-L-K-P-F-E** |
| 19 | 691 - 705 | S-I-I-A-Y-T-M-S-L-G-A-E-N-S-V |
| 20 | 1261 - 1275 | S-E-P-V-L-K-G-V-K-L-H-Y-T |
| 21 | **416 - 430** | **G-K-I-A-D-Y-N-Y-K-L-P-D-D-F-T** |
| 22 | 581 - 595 | T-L-E-I-L-D-I-T-P-C-S-F-G-G-V |
| 23 | 296 - 310 | L-S-E-T-K-C-T-L-K-S-F-T-V-E-K |
| 24 | **461 - 475** | **L-K-P-F-E-R-D-I-S-T-E-I-Y-Q-A** |
| 25 | 561 - 575 | P-F-Q-Q-F-G-R-D-I-A-D-T-T-D-A |
| 26 | 301 - 315 | C-T-L-K-S-F-T-V-E-K-G-I-Y-Q-T |
| 27 | 696 - 710 | T-M-S-L-G-A-E-N-S-V-A-Y-S-N-N |
| 28 | 1216 - 1230 | I-W-L-G-F-I-A-G-L-I-A-I-V-M-V |
| 29 | 556 - 570 | N-K-K-F-L-P-F-Q-Q-F-G-R-D-I-A |
| 30 | 1211 - 1225 | K-W-P-W-Y-I-W-L-G-F-I-A-G-L-I |

| No.3 | Position | Amino Acid Sequence |
|---|---|---|
| 1 | 816 - 830 | S-F-I-E-D-L-L-F-N-K-V-T-L-A-D |
| 2 | 691 - 705 | S-I-I-A-Y-T-M-S-L-G-A-E-N-S-V |
| 3 | 686 - 700 | S-V-A-S-Q-S-I-I-A-Y-T-M-S-L-G |
| 4 | 1176 - 1190 | V-V-N-I-Q-K-E-I-D-R-L-N-E-V-A |
| 5 | 1141- 1155 | L-Q-P-E-L-D-S-F-K-E-E-L-D-K-Y |
| 6 | 1171 - 1185 | G-I-N-A-S-V-V-N-I-Q-K-E-I-D-R |
| 7 | 1146 - 1160 | D-S-F-K-E-E-L-D-K-Y-F-K-N-H-T |
| 8 | 556 - 570 | N-K-K-F-L-P-F-Q-Q-F-G-R-D-I-A |
| 9 | 561 - 575 | P-F-Q-Q-F-G-R-D-I-A-D-T-T-D-A |
| 10 | 571 - 585 | D-T-T-D-A-V-R-D-P-Q-T-L-E-I-L |
| 11 | 681 - 695 | P-R-R-A-R-S-V-A-S-Q-S-I-I-A-Y |
| 12 | 626 - 640 | A-D-Q-L-T-P-T-W-R-V-Y-S-T-G-S |
| 13 | **311 - 325** | **G-I-Y-Q-T-S-N-F-R-V-Q-P-T-E-S** |
| 14 | 886 - 900 | W-T-F-G-A-G-A-A-L-Q-I-P-F-A-M |
| 15 | 811 - 825 | K-P-S-K-R-S-F-I-E-D-L-L-F-N-K |
| 16 | 1256 - 1270 | F-D-E-D-D-S-E-P-V-L-K-G-V-K-L |
| 17 | 696 - 710 | T-M-S-L-G-A-E-N-S-V-A-Y-S-N-N |
| 18 | 1166 - 1180 | L-G-D-I-S-G-I-N-A-S-V-V-N-I-Q |
| 19 | **321 - 335** | **Q-P-T-E-S-I-V-R-F-P-N-I-T-N-L** |
| 20 | 821 - 835 | L-L-F-N-K-V-T-L-A-D-A-G-F-I-K |
| 21 | 1151 - 1165 | E-L-D-K-Y-F-K-N-H-T-S-P-D-V-D |
| 22 | 621 - 635 | P-V-A-I-H-A-D-Q-L-T-P-T-W-R-V |
| 23 | **491 - 505** | **P-L-Q-S-Y-G-F-Q-P-T-N-G-V-G-Y** |
| 24 | 1251 - 1265 | G-S-C-C-K-F-D-E-D-D-S-E-P-V-L |
| 25 | 1131 - 1145 | E-C-V-L-G-Q-S-K-R-V-D-F-C-G-K |
| 26 | 836 - 850 | Q-Y-G-D-C-L-G-D-I-A-A-R-D-L-I |
| 27 | 176 - 190 | L-M-D-L-E-G-K-Q-G-N-F-K-N-L-R |
| 28 | 971 - 985 | G-A-I-S-S-V-L-N-D-I-L-S-R-L-D |
| 29 | 566 - 580 | G-R-D-I-A-D-T-T-D-A-V-R-D-P-Q |
| 30 | 806 - 820 | L-P-D-P-S-K-P-S-K-R-S-F-I-E-D |

| No.4 | Position | Amino Acid Sequence |
|---|---|---|
| 1 | 1146 - 1160 | D-S-F-K-E-E-L-D-K-Y-F-K-N-H-T |
| 2 | 621 - 635 | P-V-A-I-H-A-D-Q-L-T-P-T-W-R-V |
| 3 | 1131 - 1145 | G-I-V-N-N-T-V-Y-D-P-L-Q-P-E-L |
| 4 | 1016 - 1030 | A-E-I-R-A-S-A-N-L-A-A-T-K-M-S |
| 5 | 946 - 960 | G-K-L-Q-D-V-V-N-Q-N-A-Q-A-L-N |
| 6 | 1261 - 1275 | S-E-P-V-L-K-G-V-K-L-H-Y-T |
| 7 | 986 - 1000 | K-V-E-A-E-V-Q-I-D-R-L-I-T-G-R |
| 8 | 1006 - 1020 | T-Y-V-T-Q-Q-L-I-R-A-A-E-I-R-A |
| 9 | 996 - 1010 | L-I-T-G-R-L-Q-S-L-Q-T-Y-V-T-Q |
| 10 | 1266 - 1280 | E-P-V-L-K-G-V-K-L-H-Y-T |
| 11 | 851 - 865 | C-A-Q-K-F-N-G-L-T-V-L-P-P-L-L |
| 12 | 121 - 135 | N-N-A-T-N-V-V-I-K-V-C-E-F-Q-F |
| 13 | 1101 - 1115 | H-W-F-V-T-Q-R-N-F-Y-E-P-Q-I-I |
| 14 | 846 - 860 | A-R-D-L-I-C-A-Q-K-F-N-G-L-T-V |
| 15 | 1096 -1110 | V-S-N-G-T-H-W-F-V-T-Q-R-N-F-Y |
| 16 | 971 - 985 | G-A-I-S-S-V-L-N-D-I-L-S-R-L-D |
| 17 | **456 - 470** | **F-R-K-S-N-L-K-P-F-E-R-D-I-S-T** |
| 18 | 1256 - 1270 | F-D-E-D-D-S-E-P-V-L-K-G-V-K-L |
| 19 | 1 - 15 | M-F-V-F-L-V-L-L-P-L-V-S-S-Q-C |
| 20 | 1211 - 1225 | K-W-P-W-Y-I-W-L-G-F-I-A-G-L-I |
| 21 | 951 - 965 | V-V-N-Q-N-A-Q-A-L-N-T-L-V-K-Q |
| 22 | 181 - 195 | G-K-Q-G-N-F-K-N-L-R-E-F-V-F-K |
| 23 | 811 - 825 | K-P-S-K-R-S-F-I-E-D-L-L-F-N-K |
| 24 | 116 - 130 | S-L-L-I-V-N-N-A-T-N-V-V-I-K-V |
| 25 | **506 - 520** | **Q-P-Y-R-V-V-V-L-S-F-E-L-L-H-A** |
| 26 | 126 - 140 | V-V-I-K-V-C-E-F-Q-F-C-N-D-P-F |
| 27 | 231 - 245 | I-G-I-N-I-T-R-F-Q-T-L-L-A-L-H |
| 28 | 1251 - 1265 | G-S-C-C-K-F-D-E-D-D-S-E-P-V-L |
| 29 | 991 - 1005 | V-Q-I-D-R-L-I-T-G-R-L-Q-S-L-Q |
| 30 | **431 - 445** | **G-C-V-I-A-W-N-S-N-N-L-D-S-K-V** |

| No.5 | Position | Amino Acid Sequence |
|---|---|---|
| 1 | 116- 130 | S-L-L-I-V-N-N-A-T-N-V-V-I-K-V |
| 2 | 221 - 235 | S-A-L-E-P-L-V-D-L-P-I-G-I-N-I |
| 3 | 661 - 675 | E-C-D-I-P-I-G-A-G-I-C-A-S-Y-Q |
| 4 | 1211 - 1225 | K-W-P-W-Y-I-W-L-G-F-I-A-G-L-I |
| 5 | 621 - 635 | P-V-A-I-H-A-D-Q-L-T-P-T-W-R-V |
| 6 | **346 - 360** | **R-F-A-S-V-Y-A-W-N-R-K-R-I-S-N** |
| 7 | 616 - 630 | N-C-T-E-V-P-V-A-I-H-A-D-Q-L-T |
| 8 | **506 - 520** | **Q-P-Y-R-V-V-V-L-S-F-E-L-L-H-A** |
| 9 | 1221 - 1235 | I-A-G-L-I-A-I-V-M-V-T-I-M-L-C |
| 10 | 106 - 120 | F-G-T-T-L-D-S-K-T-Q-S-L-L-I-V |
| 11 | 121 - 135 | N-N-A-T-N-V-V-I-K-V-C-E-F-Q-F |
| 12 | 811 - 825 | K-P-S-K-R-S-F-I-E-D-L-L-F-N-K |
| 13 | **326 - 340** | **I-V-R-F-P-N-I-T-N-L-C-P-F-G-E** |
| 14 | 1216 - 1230 | I-W-L-G-F-I-A-G-L-I-A-I-V-M-V |
| 15 | 1121 -1135 | F-V-S-G-N-C-D-V-V-I-G-I-V-N-N |
| 16 | 851 - 865 | C-A-Q-K-F-N-G-L-T-V-L-P-P-L-L |
| 17 | 971 - 985 | G-A-I-S-S-V-L-N-D-I-L-S-R-L-D |
| 18 | **451 - 465** | **Y-L-Y-R-L-F-R-K-S-N-L-K-P-F-E** |
| 19 | 1126 - 1240 | C-D-V-V-I-G-I-V-N-N-T-V-Y-D-P |
| 20 | 76 - 95 | T-K-R-F-D-N-P-V-L-P-F-N-D-G-V |
| 21 | 1 - 15 | M-F-V-F-L-V-L-L-P-L-V-S-S-Q-C |
| 22 | **486 - 500** | **F-N-C-Y-F-P-L-Q-S-Y-G-F-Q-P-T** |
| 23 | 706 - 720 | A-Y-S-N-N-S-I-A-I-P-T-N-F-T-I |
| 24 | 946 - 960 | G-K-L-Q-D-V-V-N-Q-N-A-Q-A-L-N |
| 25 | 1056 - 1070 | A-P-H-G-V-V-F-L-H-V-T-Y-V-P-A |
| 26 | 876 - 890 | A-L-L-A-G-T-I-T-S-G-W-T-F-G-A |
| 27 | 586 - 600 | D-I-T-P-C-S-F-G-G-V-S-V-I-T-P |
| 28 | 176 - 190 | L-M-D-L-E-G-K-Q-G-N-F-K-N-L-R |
| 29 | 111 - 125 | D-S-K-T-Q-S-L-L-I-V-N-N-A-T-N |
| 30 | 846 - 860 | A-R-D-L-I-C-A-Q-K-F-N-G-L-T-V |

| No.6 | Position | Amino Acid Sequence |
|---|---|---|
| 1 | 971 - 985 | G-A-I-S-S-V-L-N-D-I-L-S-R-L-D |
| 2 | 691 - 705 | S-I-I-A-Y-T-M-S-L-G-A-E-N-S-V |
| 3 | 686 - 700 | S-V-A-S-Q-S-I-I-A-Y-T-M-S-L-G |
| 4 | 1001 - 1015 | L-Q-S-L-Q-T-Y-V-T-Q-Q-L-I-R-A |
| 5 | 76 - 95 | T-K-R-F-D-N-P-V-L-P-F-N-D-G-V |
| 6 | 946 - 960 | G-K-L-Q-D-V-V-N-Q-N-A-Q-A-L-N |
| 7 | 851 - 865 | C-A-Q-K-F-N-G-L-T-V-L-P-P-L-L |
| 8 | 116 - 130 | S-L-L-I-V-N-N-A-T-N-V-V-I-K-V |
| 9 | 1256 - 1270 | F-D-E-D-D-S-E-P-V-L-K-G-V-K-L |
| 10 | 1176 - 1190 | V-V-N-I-Q-K-E-I-D-R-L-N-E-V-A |
| 11 | 846 - 860 | A-R-D-L-I-C-A-Q-K-F-N-G-L-T-V |
| 12 | 621 - 635 | P-V-A-I-H-A-D-Q-L-T-P-T-W-R-V |
| 13 | **491 - 505** | **P-L-Q-S-Y-G-F-Q-P-T-N-G-V-G-Y** |
| 14 | 1251 - 1265 | G-S-C-C-K-F-D-E-D-D-S-E-P-V-L |
| 15 | 681 - 695 | P-R-R-A-R-S-V-A-S-Q-S-I-I-A-Y |
| 16 | 1211 - 1255 | I-A-G-L-I-A-I-V-M-V-T-I-M-L-C |
| 17 | 106 - 120 | F-G-T-T-L-D-S-K-T-Q-S-L-L-I-V |
| 18 | 1171 - 1185 | G-I-N-A-S-V-V-N-I-Q-K-E-I-D-R |
| 19 | 191 - 205 | E-F-V-F-K-N-I-D-G-Y-F-K-I-Y-S |
| 20 | 1131 - 1145 | G-I-V-N-N-T-V-Y-D-P-L-Q-P-E-L |
| 21 | **321 - 335** | **Q-P-T-E-S-I-V-R-F-P-N-I-T-N-L** |
| 22 | 1216 - 1230 | I-W-L-G-F-I-A-G-L-I-A-I-V-M-V |
| 23 | **326 - 340** | **I-V-R-F-P-N-I-T-N-L-C-P-F-G-E** |
| 24 | **431 - 445** | **G-C-V-I-A-W-N-S-N-N-L-D-S-K-V** |
| 25 | **416 - 430** | **G-K-I-A-D-Y-N-Y-K-L-P-D-D-F-T** |
| 26 | 91 - 105 | Y-F-A-S-T-E-K-S-N-I-I-R-G-W-I |
| 27 | **456 - 470** | **F-R-K-S-N-L-K-P-F-E-R-D-I-S-T** |
| 28 | 906 - 920 | F-N-G-I-G-V-T-Q-N-V-L-Y-E-N-Q |
| 29 | 221 - 235 | S-A-L-E-P-L-V-D-L-P-I-G-I-N-I |
| 30 | 136 - 150 | C-N-D-P-F-L-G-V-Y-Y-H-K-N-N-K |
